# A quantitative study of the Golgi retention of glycosyltransferases

**DOI:** 10.1101/2021.02.15.431224

**Authors:** Xiuping Sun, Bing Chen, Zhiwei Song, Lei Lu

## Abstract

How Golgi glycosyltransferases and glycosidases (hereafter glycosyltransferases) localize to the Golgi is still unclear. Here, we first investigated the post-Golgi trafficking of glycosyltransferases. We found that glycosyltransferases can escape the Golgi to the plasma membrane, where they are subsequently endocytosed to the endolysosome. Post-Golgi glycosyltransferases are probably degraded by the ecto-domain shedding. We discovered that most glycosyltransferases are not retrieved from post-Golgi sites, indicating that retention but not retrieval should be the main mechanism for their Golgi localization. We proposed to use the Golgi residence time to study the Golgi retention of glycosyltransferases quantitatively and systematically. Various chimeras between ST6GAL1 and either transferrin receptor or tumor necrosis factor α quantitatively revealed the contributions of three regions of ST6GAL1, namely the N-terminal cytosolic tail, transmembrane domain and ecto-domain, to the Golgi retention. We found that each of the three regions is sufficient to produce a retention in an additive manner. The N-terminal cytosolic tail length negatively affects the Golgi retention of ST6GAL1, similar to what is known of the transmembrane domain. Therefore, long N-terminal cytosolic tail and transmembrane domain can be a Golgi export signal for transmembrane secretory cargos.

## INTRODUCTION

In addition to serving as the membrane trafficking hub, the Golgi also carries out glycosylation modifications for secretory proteins and lipids (cargos). Golgi glycosyltransferases and glycosidases (hereafter glycosyltransferases) are the major transmembrane residents of the Golgi, which synthesize or modify diverse ranges of glycans (Stanley, 2011). Like typical transmembrane secretory cargos, glycosyltransferases are synthesized in the endoplasmic reticulum (ER) before entering the secretory pathway to the Golgi. How glycosyltransferases, but not the post-Golgi targeted transmembrane secretory cargos, are retained at the Golgi is still incompletely understood.

Glycosyltransferases are type II transmembrane proteins consisting of a short N-terminal cytosolic tail (NCT), transmembrane domain (TMD) and ecto-domain (ED). The ED comprises two parts: the juxtamembrane stem region and the membrane distal catalytic domain. Extensive mutagenesis studies suggested that all these regions are important for the Golgi localization (Banfield, 2011). Three major models have emerged to account for the Golgi localization of glycosyltransferases (Banfield, 2011). In the kin-recognition model (Nilsson et al., 1994; Nilsson et al., 1993), glycosyltransferases were proposed to assemble as large oligomers or aggregates which prevent their subsequent Golgi export. Supporting this model, many glycosyltransferases have been reported to assemble as complexes (Kellokumpu et al., 2016). The TMD-based sorting model proposes that the retention or export at the Golgi is determined by TMD length (Bretscher and Munro, 1993; Munro, 1995a; Munro, 1995b). As the thickness of membrane lipid bilayer increases along the secretory pathway from the Golgi to the plasma membrane (PM), glycosyltransferases, which have a shorter TMD than PM-targeted transmembrane cargos, are expected to better match the lipid bilayer environment of the Golgi and therefore are retained there. Recent progresses in this field also suggested that glycosyltransferases might also adopt a signal-dependent retention mechanism, in which they are retrieved from the maturing *trans*-Golgi via COPI-coated vesicles (Arakel and Schwappach, 2018; Popoff et al., 2011). The NCTs of several glycosyltransferases were found to directly bind to δ and ζ-COP (subunits of COPI coat)(Liu et al., 2018), while others seem to depend on GOLPH3 (Vps74p in yeast) to indirectly interact with COPI (Ali et al., 2012; Eckert et al., 2014; Pereira et al., 2014; Schmitz et al., 2008; Tu et al., 2008).

Despite progresses made in this field, we still lack a comprehensive understanding of the subcellular localization of glycosyltransferases. For example, it is unclear how they stay at a particular sub-Golgi location under the cisternal progression or the stable compartment models, two major views on the intra-Golgi trafficking (Glick and Luini, 2011), and how they primarily reside in the interior of the Golgi stack in contrast to the peripheral distribution of trafficking machinery components (Tie et al., 2018). Although predominantly at the Golgi, 10 - 30% of a glycosyltransferase is present at the ER due to their retrograde trafficking from the Golgi to the ER (Cole et al., 1998; Cole et al., 1996; Rhee et al., 2005; Storrie et al., 1998; Zaal et al., 1999). Small amount of endogenous and overexpressed glycosyltransferases such as B4GALT1 (GalT) and ST6GAL1 (ST) were also detected at the PM (Berger, 2002; Chen et al., 2000; Teasdale et al., 1994; Wong et al., 1992). In contrast, much less is known about their post-Golgi trafficking, probably due to their low abundance there. Since a protein’s Golgi localization is determined by retention and/or retrieval (Munro, 1998), it is important to address whether glycosyltransferases can be retrieved from post-Golgi locations, which is currently unclear. Past studies on the Golgi localization of glycosyltransferases mainly focused on their Golgi retention and much less attention has been paid to their potential retrieval from post-Golgi sites. At last, although extensive data have been collected for this topic, they were primarily qualitative descriptions. Here, by mainly using ST, we demonstrated that most glycosyltransferases are not retrieved to the Golgi and, instead, they shed their EDs extracellularly. Due to the lack of the retrieval pathway, we proposed to adopt the Golgi residence time (Sun et al., 2020) as a metric for the Golgi retention of glycosyltransferases. By this approach, we quantitatively and systematically assessed the contribution of the NCT, TMD and ED in the Golgi localization of ST. We found that all three regions additively contribute to its efficient Golgi retention while its NCT length plays a negative role.

## RESULTS

### Most glycosyltransferases are not retrieved from post-Golgi sites

From our quantitative sub-Golgi localization study (GLIM) (Tie et al., 2016), Golgi glycosyltransferases mainly reside at the medial to *trans*-Golgi but not the TGN (Tie et al., 2018). It is known that the endolysosome-targeting of a secretory cargo depends on the signal that is recognized by the TGN-localized clathrin machinery (Bonifacino and Traub, 2003; De Matteis and Luini, 2008; Guo et al., 2014). Since such signal has not been reported for glycosyltransferases, we hypothesized that, once exiting the Golgi, glycosyltransferases should follow the constitutive secretory pathway to the PM, like interleukin 2 receptor α subunit (Tac) and vesicular stomatitis virus glycoprotein G (VSVG). However, unlike Tac and VSVG, the PM-localized glycosyltransferase is invisible under the fluorescence microscopy under the overexpression. To investigate the PM presence of glycosyltransferases, live HeLa cells expressing GFP-tagged glycosyltransferases and transferrin receptor (TfR) were incubated with VHH-mCherry, a recombinant anti-GFP nanobody (Buser et al., 2018) which selectively binds to extracellularly exposed GFP. It was found that VHH-mCherry was internalized in cells expressing GFP-tagged TfR (positive control), ST and MGAT1 but not those expressing cytosolic GFP (negative control) (Fig. 1A), therefore confirming the PM localization of glycosyltransferases. We next followed the endocytic trafficking of various glycosyltransferases and asked if they can be retrieved to the Golgi. In cells expressing ST-GFP, the majority of internalized VHH-mCherry was first observed at the early endosome as marked by EEA1 after 10-20 min (Fig. 1B) and to the late endosome and lysosome at 60-120 min post internalization (Fig. 1C). With up to 8 h continuous internalization, we did not detect the localization of VHH-mCherry at the Golgi in positive internalization cells (n = 106) (first row; Fig. 1D), demonstrating that ST is likely not retrieved from the PM and endosome to the Golgi.

**Figure 1.**
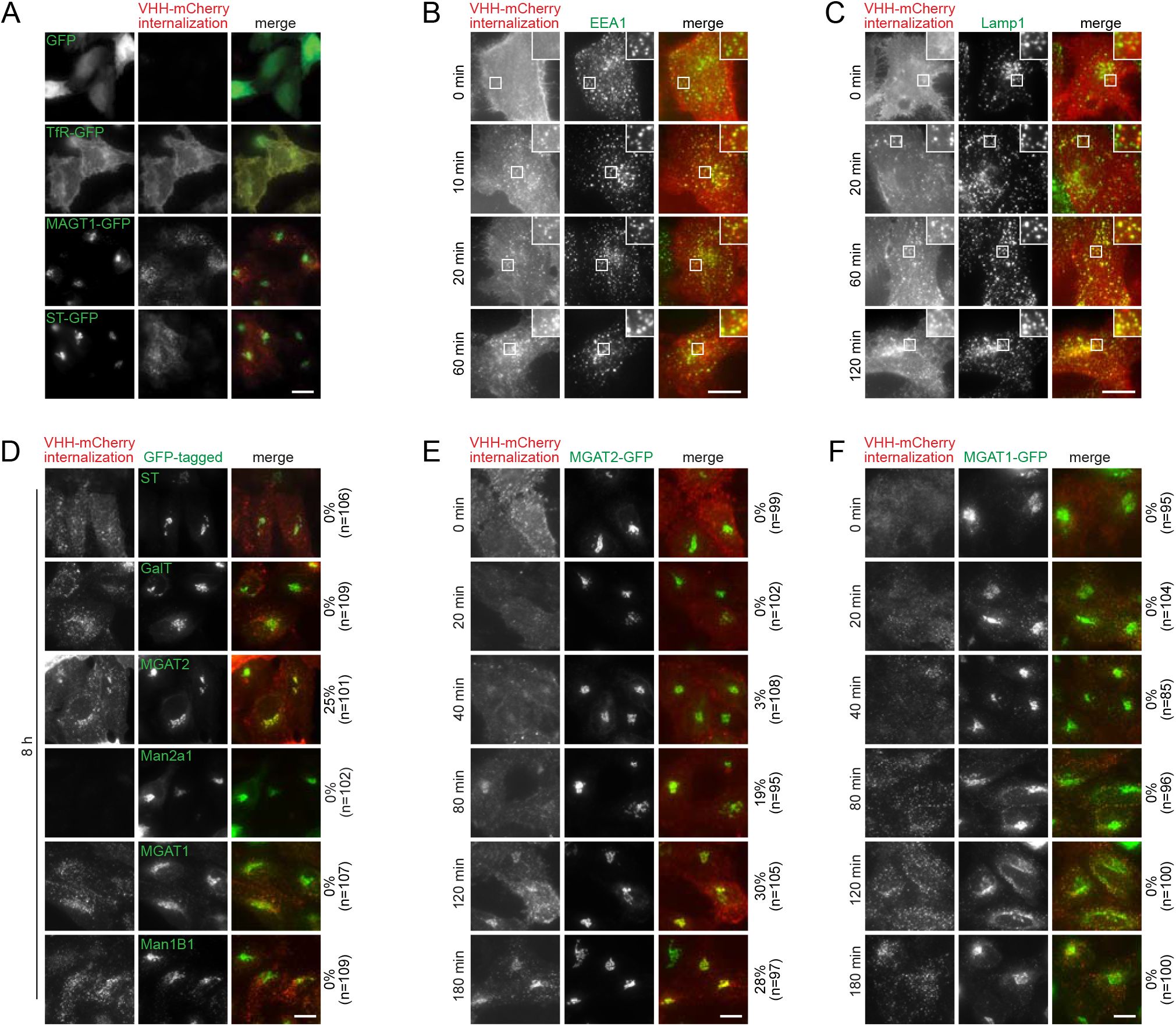
The endocytic trafficking of cell surface Golgi glycosyltransferases. (A) Cell surface MAGT1 and ST can be surface-labeled and internalized. HeLa cells transiently expressing GFP (negative control) or indicated C-terminally GFP-tagged transmembrane proteins were continuously incubated with VHH-mCherry at 37 °C for 1 h. TfR-GFP serves as a positive control. (B,C) Cell surface ST-GFP is endocytosed to the early endosome before reaching the late endosome and lysosome. HeLa cells transiently expressing ST-GFP were surface-labeled by VHH-mCherry on ice. After washing, cells were warmed up to 37 °C for indicated time and immuno-stained for endogenous EEA1 (B) and Lamp1 (C). Boxed regions are amplified in the upper right corner. (D) Among N-glycan modification glycosyltransferases, Man1B1, MGAT1, Man2a1, MGAT2, GalT and ST, only MGAT2 can be retrieved from the cell surface to the Golgi. HeLa cells transiently expressing indicated glycosyltransferases were continuously incubated with VHH-mCherry at 37 °C for 8 h. Note that the surface pool of Man2a1-GFP was undetectable by our assay. (E) The retrieval kinetics of cell surface MGAT2 to the Golgi. HeLa cells transiently expressing MGAT2-GFP were surface-labeled by VHH-mCherry on ice. After washing, cells were warmed up to 37 °C for indicated time. (F) Cell surface MGAT1 is not retrieved to the Golgi. The experiment was performed in parallel to (E) and serves as a negative control. Scale bar, 20 μm. In (D-F), the number on the right of each panel indicates the percentage of cells displaying the Golgi localization of VHH-mCherry; n, the number of cells counted.

We screened 14 additional Golgi glycosyltransferases including GFP-tagged Man1B1, MGAT1, Man2a1, MGAT2, GalT, GCNT1, GALNT4, GALNT8, TPST1 and TPST2 and Myc-tagged B3galt6, B4GALT3, B4GALT7 and POMGNT1 (Fig. 1D; Fig. S1, A and B), using VHH-mCherry (for GFP-tagged enzymes) or anti-Myc antibody (for Myc-tagged enzymes) continuous internalization assay. During 8 h’s internalization, Man2a1-GFP and GALNT8-GFP were negative in labeling, demonstrating that they do not have a detectable PM-pool (Fig. 1D; Fig. S1B). For the rest, we found that only MGAT2, B3galt6, B4GALT7 and POMGNT1 expressing cells, displayed the Golgi targeting. However, the staining pattern was heterogeneous as the Golgi localization was only found in 20 - 50% of internalization positive cells. In summary, we conclude that the majority of Golgi glycosyltransferases (11 out of 15) are not retrieved from post-Golgi sites including the endosome and PM.

The time course of VHH-mCherry internalization revealed that MGAT2-GFP was first detected at the Golgi after 40 min of continuous internalization (Fig. 1E). Longer incubation time led to up to 30% Golgi-positive cells. The Golgi targeting of the surface-labeled MGAT2-GFP was also confirmed by anti-GFP antibody (Fig. S1C). In contrast, VHH-mCherry was always negative at the Golgi in cells expressing MGAT1-GFP (Fig. 1F). The Golgi retrieval or retrograde transport of MGAT2 seems to be much slower than that of the typical retrograde cargos such as TGN38, furin and shiga toxin B fragment, which take 20 - 30 min (Ghosh et al., 1998; Mallard et al., 1998; Mallet and Maxfield, 1999). The slow kinetics and low efficiency of Golgi retrieval imply that MGAT2, B3galt6, B4GALT7 and POMGNT1 probably adopt alternative pathways and machinery, which await further elucidation. Altogether, our data suggest that 1) glycosyltransferases can escape the Golgi and reach the PM; 2) at the PM, they undergo endocytosis to sequentially transit the early endosome, late endosome and lysosome; and 3) most glycosyltransferases do not possess a post-Golgi retrieval pathway to return to the Golgi.

### Post-Golgi ST are mostly degraded by the ED-shedding

Glycosyltransferases have been reported to undergo ED-shedding to release their EDs to the extracellular space (Ohtsubo and Marth, 2006; Paulson and Colley, 1989). We analyzed the conditioned cell culture media and lysates of cells expressing ST- or MGAT1-GFP. Both ST- and MGAT1-GFP were detected in culture media as fragments that were smaller than corresponding ones in lysates (Fig. 2A; Fig. S2A), consistent with the ED-shedding.

**Figure 2.**
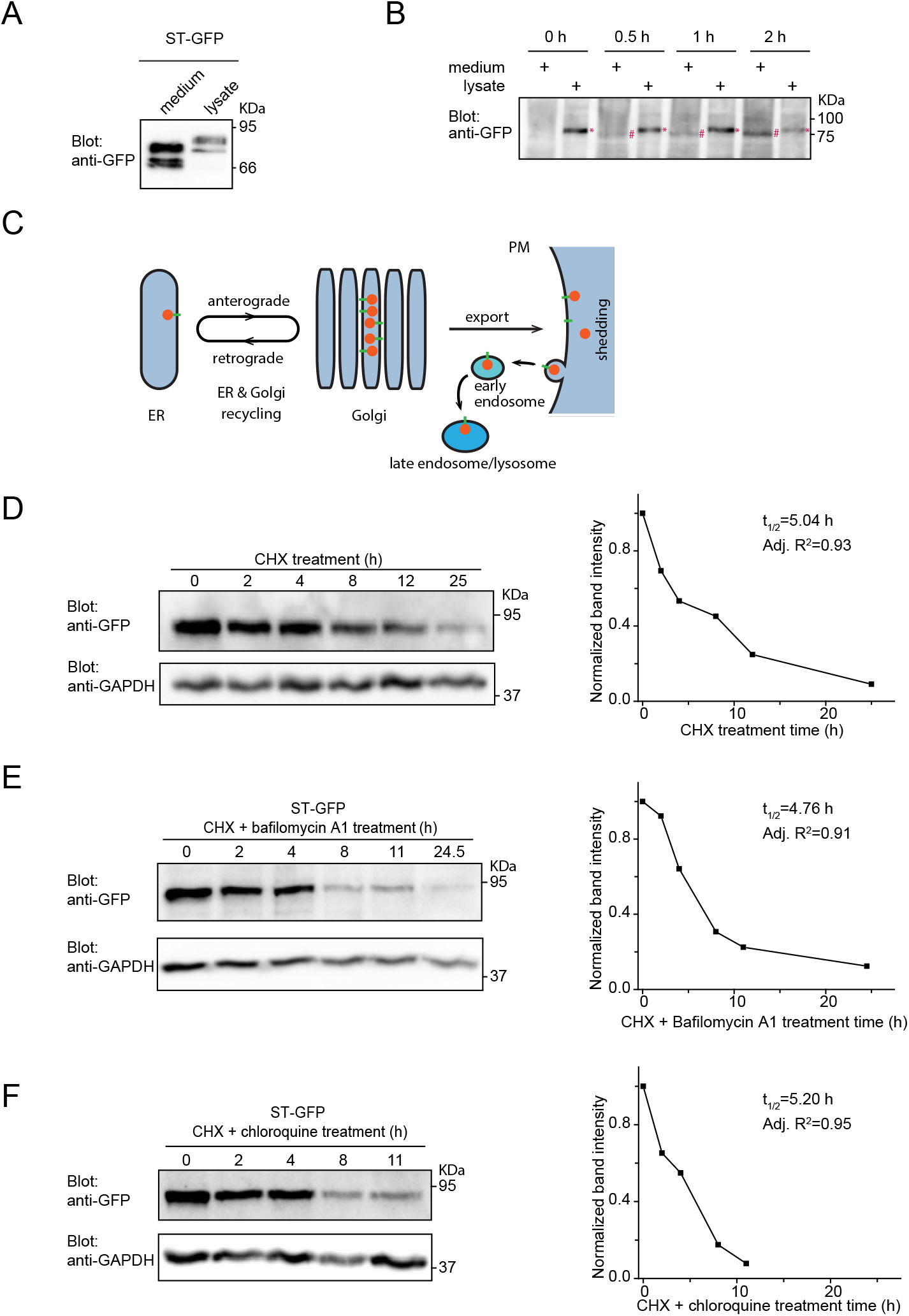
The ED-shedding of post-Golgi ST. (A) The ED of ST can be detected in the cell culture medium. HeLa cells transiently expressing ST-GFP were cultured for 24 h. The culture medium and cell lysate were subjected to Western blot analysis using anti-GFP antibody. (B) The ED-shedding kinetics of surface-labeled ST. HeLa cells transiently expressing ST-GFP were surface-labeled with VHH-mCherry on ice. After washing, cells were incubated at 37 °C for indicated time. The culture medium and corresponding cell lysate were subsequently subjected to immunoprecipitation using rabbit anti-mCherry polyclonal antibody. Pulldowns were analyzed in Western blot analysis using anti-GFP antibody. * and # indicate the full length (upper band) and ED (lower band) of ST, respectively. (C) The schematic diagram showing the intracellular trafficking itinerary of ST. See text for details. (D-F) The half-life of cellular ST is not substantially affected when the lysosomal degradation is inhibited. HeLa cells transiently expressing ST-GFP were treated with CHX (D) or CHX in combination with either bafilomycin A1 (E) or chloroquine (F) for indicated time. Cell lysates were subjected to Western blot analysis by anti-GFP or anti-GAPDH (loading control) antibody. The quantification of the gel blot was shown in the right plot. Band intensities were measured and normalized to that of 0 h. t1/2s were acquired by fitting to the first order exponential decay function. In (A, B, D-F), molecular weights (in KDa) are labeled at right in all blots.

It has been suggested that the shedding of ST can be mediated by Cathepsin D-like proteases (Lammers and Jamieson, 1988) and beta-site APP cleaving enzyme 1 (BACE1) (Kitazume et al., 2001), both having a PM and endosomal pool (Haass et al., 2012; Stoka et al., 2016). Based on the observation that 20 °C incubation blocks the cleavage and secretion of the ED, Ma *et al.* proposed that the cleavage or shedding occurs at a post-Golgi localization (Ma et al., 1997). Although 20 °C incubation arrests cargo exit from the Golgi (Saraste and Kuismanen, 1984), the possibility that it also inhibits the cleavage of ST was not ruled out in their study. We conducted the below experiment to investigate if ED-shedding can occur at the post-Golgi localization. The cell surface pool of ST-GFP was first selectively labeled by VHH-mCherry on ice. After washing unbound VHH-mCherry, cells were warmed up to 37 °C for various length of time before collecting the culture medium and cell lysate. At last, the VHH-mCherry-labeled ST-GFP was immunoprecipitated by anti-mCherry antibody and analyzed in Western blot (Fig. 2B). We found that the ED of surface-labeled ST was detected in the medium 0.5 h after warming up to 37 °C and most surface-labeled ST was cleaved within 2 h. Since surface ST is not retrieved to the Golgi (see above), our data demonstrate that ED-shedding of ST probably occurs at the PM. Combining with our new data, the intracellular trafficking itinerary of ST is illustrated in Fig. 2C. Note that, at the moment, we cannot rule out the possibility that the cleavage takes place at the endolysosome, which subsequently undergoes exocytosis to release the ED (Luzio et al., 2007).

Under the cycloheximide (CHX) treatment, we observed that the total cellular ST gradually disappeared with a half-life of 5.0 h (Fig. 2D). The similarity between the degradation half-life and the Golgi residence time of ST (~ 5h; see below) suggests that the Golgi exocytic export to the PM is probably the rate-limiting step. We found that the half-lives of ST-GFP remained roughly the same as the control in the presence of the lysosomal degradation inhibitor, bafilomycin A1 or chloroquine (Fig. 2, E and F), suggesting that the ED-shedding but not the lysosomal degradation might be the major post-Golgi turnover pathway for ST.

### The Golgi residence time is a quantitative metric of the Golgi retention

Without the post-Golgi retrieval, the Golgi localization of most glycosyltransferases is hence primarily contributed by their retention at the Golgi. Like a constitutive secretory cargo such as Tac and VSVG, the Golgi pool of a glycosyltransferase is determined by the net effect of the following three trafficking pathways, as we previously discussed (Fig. 3A) (Sun et al., 2020): 1) the ER-to-Golgi (anterograde), 2) the Golgi-to-ER (retrograde) and 3) the Golgi-to-PM (exocytic) trafficking pathways. The ER-to-Golgi transport increases the Golgi pool while the Golgi-to-PM and Golgi-to-ER pathways deplete it. For a glycosyltransferase, since the cycling between the ER and Golgi is relatively fast in comparison to the Golgi export (Rhee et al., 2005; Zaal et al., 1999), the cycling can be considered at the steady state and its net contribution to the Golgi pool is the biosynthesis from the ER. When the protein synthesis is stopped by CHX treatment, the Golgi pool of the glycosyltransferase should gradually reduce by following the first order exponential decay. Previously, we defined the Golgi residence time as the half-life, *t_1/2_*, of the Golgi pool’s fluorescence decay and used it to describe the Golgi export (Sun et al., 2020). We reasoned that the Golgi retention should be inversely related to the Golgi export — a strong Golgi export can be viewed as a weak Golgi retention and vice versa. We hence proposed the usage of the Golgi residence time as a quantitative metric for the Golgi retention.

**Figure 3.**
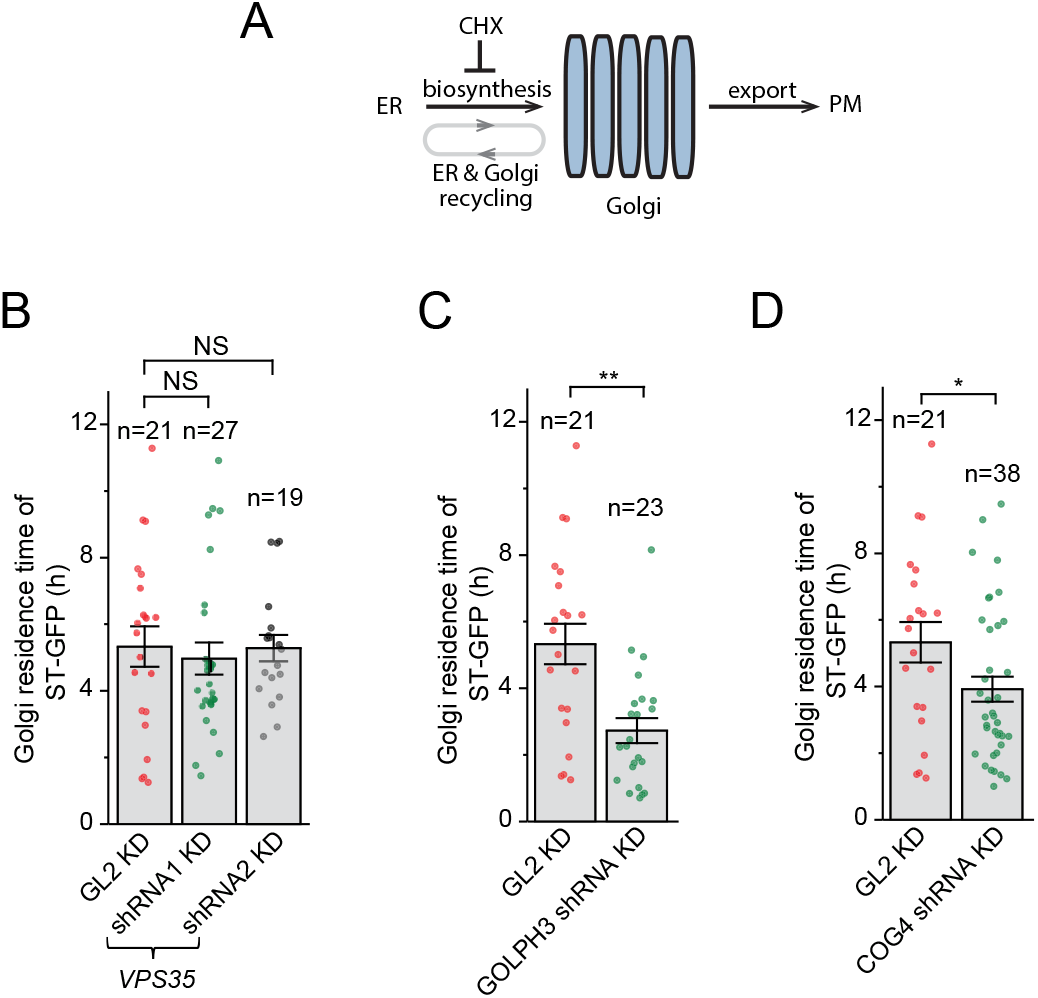
The Golgi residence time as a metric for the Golgi retention. (A) The schematic diagram illustrating trafficking pathways that contribute to the Golgi localization of a glycosyltransferase. See text for details. (B-D) The Golgi residence time of ST is substantially reduced when GOLPH3 and COG complex, but not retromer complex, are compromised. HeLa cells were subjected to lentivirus-mediated transduction of shRNAs targeting *VPS35* (a subunit of retromer complex), *GOLPH3* or *COG4* (a subunit of COG complex). Cells were subsequently transfected to express ST-GFP and the Golgi residence times were acquired (see also Fig. S3, D-F). GL2 shRNA is a non-targeting negative control. Error bar, mean ± standard error; *P* values are from *t* test (unpaired and two-tailed); NS, not significant; *, *P* ≤ 0.05; **, *P* ≤ 0.005; n, the number of quantified cells.

To validate this concept, we measured the Golgi residence time of ST when its Golgi retention is compromised. The conserved oligomeric Golgi (COG) complex (Blackburn et al., 2018) and GOLPH3 (Eckert et al., 2014; Schmitz et al., 2008; Tu et al., 2008) have been documented to regulate the retention of glycosyltransferases such as ST. Retromer complex, which functions in the carrier biogenesis at the endosome during the endosome-to-Golgi trafficking (Chen et al., 2019; Lu and Hong, 2014; McNally and Cullen, 2018), was selected as a control. The depletion of COG, GOLPH3 and retromer was achieved by lentivirus-transduced shRNAs that target to *COG4* (a subunit of COG), *GOLPH3* and *Vps35* (a subunit of retromer), respectively. Our quantitative reverse transcription PCR (RT-qPCR) confirmed that their transcripts were reduced by ≥ 78 % (Fig. S3, A-C). We found that the Golgi residence time of ST in *VPS35* knockdown cells (≥ 5.0 h) was similar to that of GL2 non-target control (5.3 h) (Fig. 3, B; Fig. S3, D). In contrast, the Golgi residence times of ST in *COG* and *GOLPH3* knockdown cells reduced substantially to 2.7 and 3.9 h, respectively (Fig. 3, C and D; Fig. S3, E and F). Our observation is therefore consistent with what we know of the Golgi retention by COG and GOLPH3, hence validating the Golgi residence time as a metric for the Golgi retention.

### The Golgi retention of ST is additively contributed by its NCT, TMD and ED

To further understand the mechanism behind the Golgi retention of a glycosyltransferase, we compared ST with TfR and tumor necrosis factor α (TNFα), two PM-targeted cargos. All three are type II transmembrane proteins comprising the NCT, TMD and ED (Fig. 4, A and B). The Golgi residence times of TfR and TNFα, < 14 min, are similar to other constitutive transmembrane secretory cargos such as Tac and VSVG and their Golgi localization is undetectable at the steady state (Sun et al., 2020). In contrast, the Golgi residence time of ST, ~ 5 h, is among the longest. Therefore, it is reasonable to assume that TfR and TNFα have no intrinsic Golgi retention. We quantitatively studied the role of the NCT, TMD and ED in the Golgi retention of ST by swapping individual region(s) with corresponding one(s) of TfR or TNFα. A series of GFP-tagged swapping chimeras were hence generated (Fig. 4, A and B) and named after the source of the NCT, TMD and ED using S, F and N, which denote ST, TfR and TNFα, respectively. For example, SFF is the chimera with the NCT from ST but the TMD and ED from TfR. By this notion, ST, TfR and TNFα are also named SSS, FFF and NNN, respectively.

**Figure 4.**
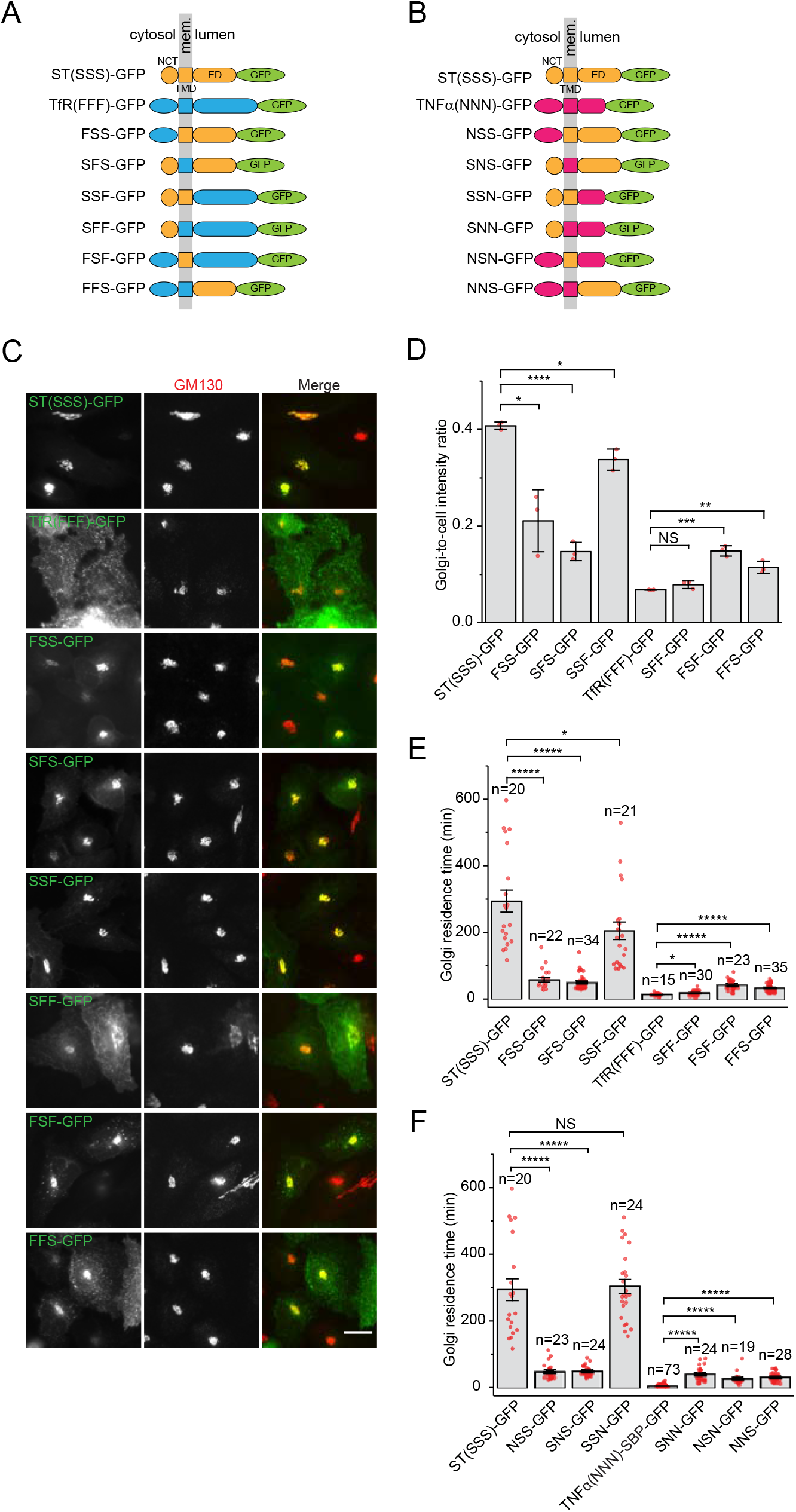
The Golgi retention of ST is contributed by its NCT, TMD and ED. (A,B) The schematic diagram showing the domain organization of swapping chimeras between ST and TfR or TNFα. Mem., membrane. (C) The subcellular localization of swapping chimeras between ST and TfR. HeLa cells transiently expressing indicated ST chimeras were immuno-stained for endogenous GM130. Scale bar, 20 μm. (D) The Golgi-to-cell intensity ratios of swapping chimeras between ST and TfR. Quantification was performed on cells described in (C). Error bar, mean ± standard deviation; n= 3 independent experiments were conducted with ≥ 20 cells quantified in each experiment. (E,F) The Golgi residence times of swapping chimeras. HeLa cells transiently expressing indicated GFP-tagged chimera and GalT-mCherry were treated with CHX and the Golgi residence times were subsequently acquired by live cell imaging (see also Fig. S4, G-T). In (F), the residence time of TNFα was from our previous study (Sun et al., 2020). Error bar, mean ± standard error (E and F). In (D-F), *P* values are from *t* test (unpaired and two-tailed); NS, not significant; *, *P* ≤ 0.05; **, *P* ≤ 0.005; ***, *P* ≤ 0.0005; ****, *P* ≤ 0.00005; *****, *P* ≤ 0.000005; n, the number of quantified cells.

Western blots demonstrated that all chimeras had expected molecular weights when expressed in cells (Fig. S4, A and B). Under the fluorescence microscopy, they displayed different extent of Golgi and PM localization (revealed by the surface staining) (Fig. 4C; Fig. S4, C-E). Therefore, all chimeras appeared to embed themselves in the membrane with their C-termini exposed to the lumen or extracellular space, consistent with their type II transmembrane topology. Their Golgi localization was quantified by the

Golgi-to-cell intensity ratio (Fig. 4D; Fig. S4F). Through the live cell imaging, the Golgi residence times of all swapping chimeras were acquired and found to be less than that of ST (Fig. 4, E and F; Fig. S4, G-T), implying that each of the three regions is likely required for the efficient Golgi retention. When more regions of ST are changed to those of TfR or TNFα, the resulting chimeras had less Golgi residence times, suggesting that the effect of the three regions on the Golgi retention might be additive for ST. On the other hand, when a region of TfR or TNFα was singly swapped with the corresponding one of ST, the resulting chimeras, SFF, FSF, FFS, SNN, NSN and NNS, had higher Golgi-to-cell intensity ratios and longer Golgi residence times than TfR or TNFα (Fig. 4, D-F; Fig. S4F), implying that each of the three regions might be sufficient to provide the Golgi retention. The Golgi residence times of these single-swapping chimeras are < 1/6 that of ST (~ 5 h) (Fig. 4, E and F), further suggesting that the Golgi retention contributed by each region might be relatively weak. In Figure 4E, while SFF (19 min) and FSF (43 min) have ≤ 2-fold increase in their Golgi residence times in comparison to TfR (14 min), SSF (204 min) displays a disproportionate 13.6-fold increase. Similar trend was also found in TNFα series of chimeras (Fig. 4F). These observations hence suggest a non-linear or synergistic effect of combining ST’s NCT and TMD. Supporting this notion, it was also observed that, while the Golgi residence times of SSF and SSN are slightly shorter than that of ST, those of FSS, SFS, NSS and SNS were < 1/5 that of ST (Fig. 4, E and F).

Interestingly, for each chimera, the Golgi-to-cell intensity ratio showed similar trend as the Golgi residence time, indicating that both metrics reflect the Golgi retention. However, the Golgi-to-cell intensity ratio is also expected to depend on a glycosyltransferase’s presence at extra-Golgi pools such as the ER, PM and endolysosome, which can be subject to degradation. Hence, the Golgi residence time is a preferred metric for the Golgi retention. Together, our data demonstrate that the Golgi retention of a glycosyltransferase is probably additively contributed by its NCT, TMD and ED.

### The NCT length negatively affects the Golgi retention of ST

We subsequently examined each of the three regions for its role in the Golgi retention of ST. We noted that glycosyltransferases tend to have a short NCT; in contrast, PM-targeted type II transmembrane proteins seem to have NCTs of diverse lengths. For example, the NCTs of ST, MAGT1 and MGAT2 contain 9, 4 and 9 amino acids (AAs) while those of TfR and TNFα contain 61 and 29 AAs, respectively. We were wondering if the NCT length plays a role in the Golgi retention. To test this hypothesis, streptavidin binding peptide or SBP (38 AAs) and FK506 binding protein or FKBP (108 AAs) were appended to the N-terminus of ST* (a clone that has truncation in the ED) to construct chimeras with longer NCTs (Fig. 5A). After confirming these chimeras (Fig. S5, A and B), their Golgi residence times (Fig. 5B; Fig. S5, C-E) showed that increasing the NCT length decreases the Golgi retention of ST*.

**Figure 5.**
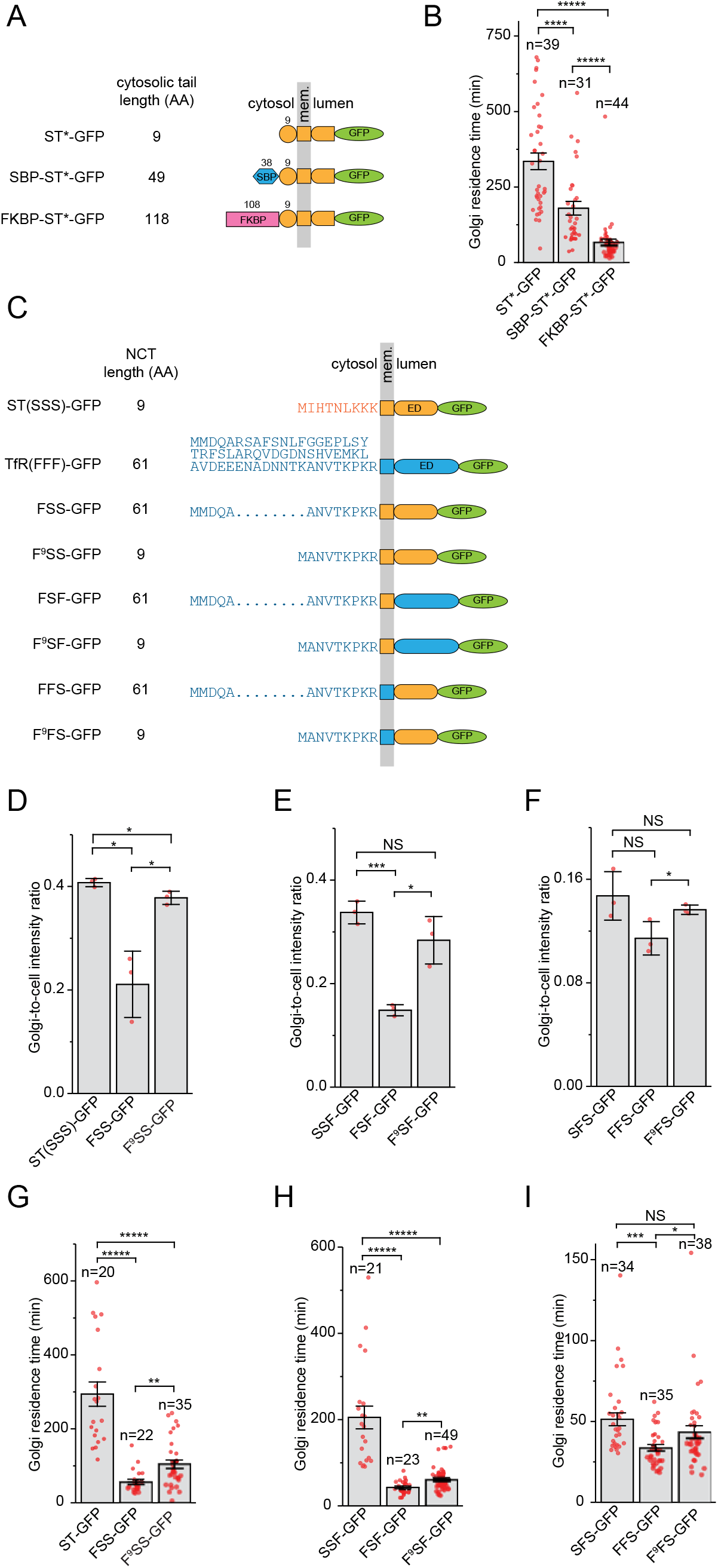
The NCT length negatively regulates the Golgi retention of ST. (A) The schematic diagram showing the domain organization of N-terminally SBP and FKBP-fused ST*. ST* has a truncated ED. Mem., membrane. (B) The Golgi residence times of chimeras in (A). HeLa cells transiently expressing GFP-tagged ST* chimeras and GalT-mCherry were treated with CHX and the Golgi residence times were subsequently acquired by live cell imaging (see also Fig. S5, C-E). (C) The schematic diagram showing the domain organization of ST and TfR swapping chimeras with different NCT lengths. (D-F) The Golgi-to-cell intensity ratios of chimeras shown in (C). HeLa cells transiently expressing indicated chimeras were immuno-stained for endogenous GM130 and the Golgi-to-cell intensity ratios were subsequently quantified. See also Figure S5H. Error bar, mean ± standard deviation; n= 3 independent experiments were conducted with ≥ 22 cells quantified in each experiment. (G-I) The Golgi residence times of chimeras shown in (C). The experiment was conducted as in (B). See also Figure S5, I-K. In (B, G-I), error bar, mean ± standard error. In (B, D-I), *P* values are from *t* test (unpaired and two-tailed); NS, not significant; *, *P* ≤ 0.05; **, *P* ≤ 0.005; ***, *P* ≤ 0.0005; ****, *P* ≤ 0.00005; *****, *P* ≤ 0.000005; n, the number of quantified cells.

Next, we tested the effect of reducing the NCT length on the Golgi retention of FSS, FSF and FFS, which all possess the NCT of TfR (Fig. 4A). By N-terminal truncation, their NCTs were subsequently shortened from 61 to 9 AAs, the same length as that of ST, to make chimeras F^9^SS, F^9^SF and F^9^FS, respectively (Fig. 5C; Fig. S5, F and G). It was found that the truncation increased their Golgi-to-cell intensity ratios (Fig. 5, D-F; Fig. S5H) and Golgi residence times (Fig. 5, G-I; Fig. S5, I-K), suggesting that increasing the NCT length might reduce the Golgi retention. In addition to the identical TMD and ED, F^9^SS and ST also have the same NCT length but with distinct sequences (Fig. 5C). However, the Golgi residence time of F^9^SS is about 1/3 that of ST (Fig. 5G). The similar trend can also be observed in F^9^SF (Fig. 5H), consistent with the synergistic effect between the NCT and TMD of ST (see above). In summary, our quantitative data demonstrate that the NCT length negatively affects the Golgi retention, therefore suggesting that the long NCT of PM-targeted type II transmembrane proteins might function as a Golgi export signal.

### Long TMD reduces the Golgi retention of ST

The TMD length has been known as a determining factor for a protein’s localization along the secretory pathway (Bretscher and Munro, 1993; Munro, 1995a; Munro, 1995b). Generally, the TMD lengths of PM membrane proteins are longer than those of Golgi ones with average of 24.4 and 20.6 AAs, respectively (Sharpe et al., 2010). For example, the TMD length of TfR is 27 AAs while that of ST is 17 AAs. Using ST*, we prepared a series of TMD insertion mutants, namely TMD18, 19, 20, 21, 22 and 24, which have increasing TMD lengths from 18 to 24 AAs (Fig. 6A). To be consistent with the previous study (Munro, 1995b), a combination of AAs, including Ala, Val and Leu, were inserted at similar position within the TMD. All mutants expressed in cells showed proteins of expected sizes and displayed extracellularly exposed C-termini (Fig. S6, A and B). TMD18-24 were observed to localize to the Golgi although the Golgi localization appeared to become poorer with the increase of the TMD (Fig. S6C). The Golgi residence times of WT and TMD18-24 revealed that the Golgi residence time decreases with the increase of the TMD length (Fig. 6B; Fig. S6, D-J). By the first order exponential decay fitting, the Golgi residence time decreases by half at the TMD length of 19.4 AAs (Fig. 6B).

**Figure 6.**
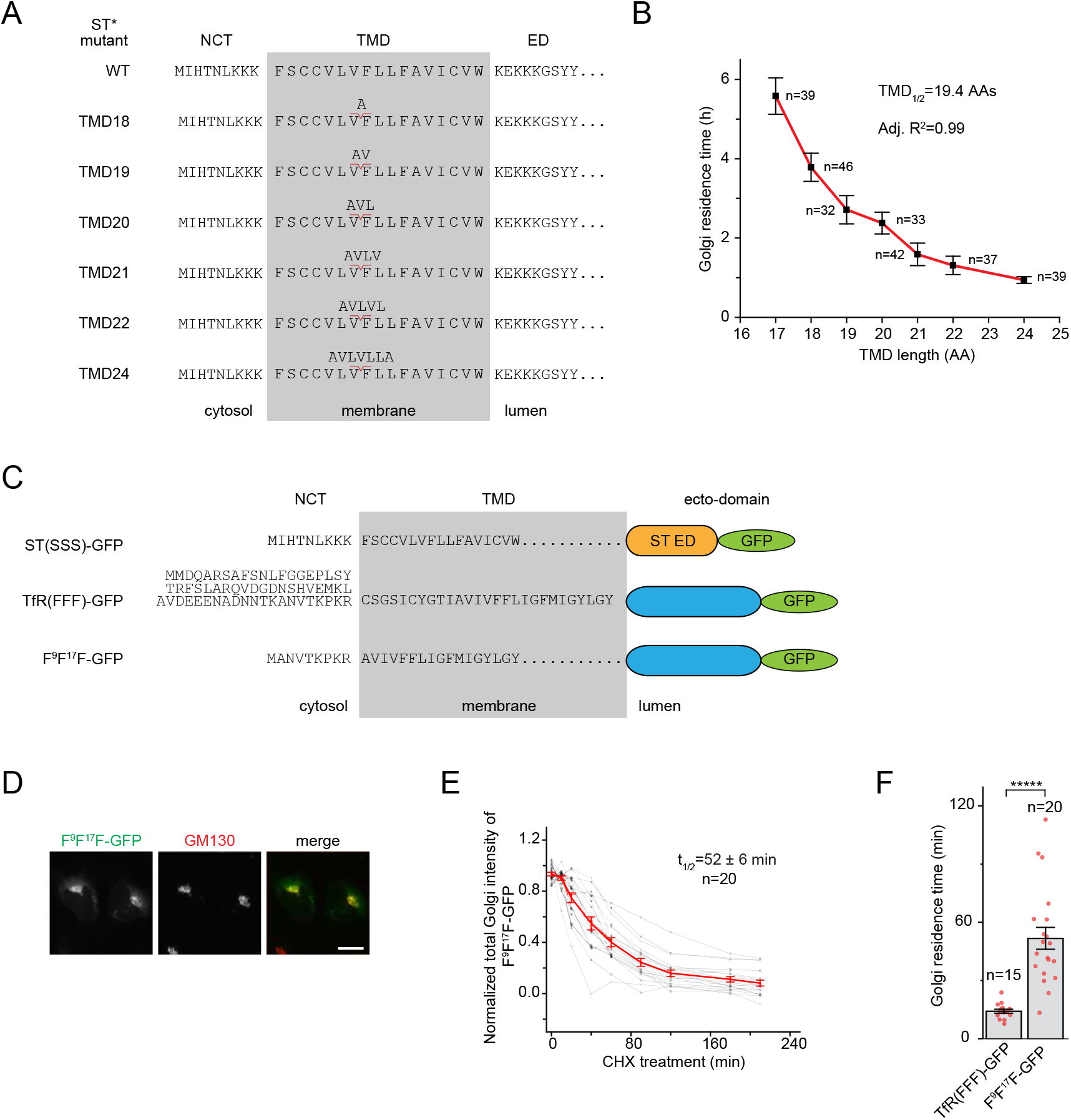
The TMD length negatively regulates the Golgi residence time of ST. (A) AA sequences of the NCT and TMD of ST* mutants with increasing TMD lengths. Inserted AAs within the TMD are indicated above. (B) The plot showing the Golgi residence time of ST* mutant vs its TMD length. HeLa cells transiently expressing GFP-tagged TMD mutant of ST* and GalT-mCherry were treated with CHX and the Golgi residence times were subsequently acquired by live cell imaging (see also Fig. S6, D-J). (C) The schematic diagram showing the NCT and TMD sequences of F^9^F^17^F in comparison to those of ST and TfR. (D) F^9^F^17^F-GFP localizes to the Golgi. HeLa cells transiently expressing F^9^F^17^F-GFP were immunostained for endogenous GM130. Scale bar, 20 μm. (E) The plot showing the Golgi intensity of F^9^F^17^F-GFP vs time. HeLa cells transiently expressing F^9^F^17^F-GFP and GalT-mCherry were subjected to live cell imaging in the presence of CHX. The data were processed as described in Figure S4, G-T. (F) The Golgi residence times of TfR-GFP and F^9^F^17^F-GFP. The Golgi residence time of F^9^F^17^F-GFP is from (E) while that of TfR-GFP was acquired similar to (E). Error bar, mean ± standard error; *P* values are from *t* test (unpaired and two-tailed); *****, *P* ≤ 0.000005; n, the number of quantified cells.

Next, we tested the effect of shortening both the TMD and NCT length on the Golgi retention of TfR. To that end, we constructed a mutant TfR, F^9^F^17^F, which has 9 and 17 AAs in the NCT and TMD respectively by truncating 52 AAs from the N-terminus and 10 AAs from the N-terminus of the TMD (Fig. 6C). Similar to other chimeras, F^9^F^17^F was confirmed by Western blot and surface staining (Fig. S6, K and L). When expressed, F^9^F^17^F-GFP predominantly localized to the Golgi (Fig. 6D), in contrast to TfR (Fig. 4C), and its Golgi residence time increased to more than 3-fold that of TfR (Fig. 6, E and F). The NCT and TMD of F^9^F^17^F have the same length as those of SSF (Fig. 4E), but its Golgi residence time is about a quarter of SSF’s, demonstrating that, in addition to sizes, their sequences probably play a more critical role in the Golgi retention, further supporting their synergistic action (see above). In summary, our data quantitatively demonstrate that the TMD length negatively regulates the Golgi retention of ST, therefore suggesting that the long TMD of PM-targeted type II transmembrane proteins might function as a Golgi export signal.

### The ED of ST is sufficient for the Golgi retention

In comparison to ST, the substantially reduced Golgi residence time of our ED-swapping chimera, SSF, demonstrates that the ED is essential for the efficient Golgi retention of ST (Fig. 4E). On the other hand, compared to TfR and TNFα, increased Golgi residence times of FFS and NNS indicate that the ED of ST can provide a Golgi retention (Fig. 4, E and F). To further investigate the role of the ED in the Golgi retention, we constructed a mCherry-tagged and GPI-anchored ED of ST, ED-mCherry-GPI (Fig. 7A). The resulting chimera is expected to be membrane attached and have the ED (together with fused mCherry) exposed to the lumen, resembling ST. The chimera and the corresponding control, SBP-mCherry-GPI, expressed proteins of expected sizes and displayed extracellular exposed mCherry, consistent with their design. In the subsequent localization studies, we found that ED-mCherry-GPI, but not the control SBP-mCherry-GPI, localized to the Golgi and also had a much longer Golgi residence time than the latter (46 vs 17 min) (Fig. 7, B-D)(Sun et al., 2020). Our results hence demonstrate that the ED of ST and possibly other glycosyltransferases can contribute to their Golgi retention.

**Figure 7.**
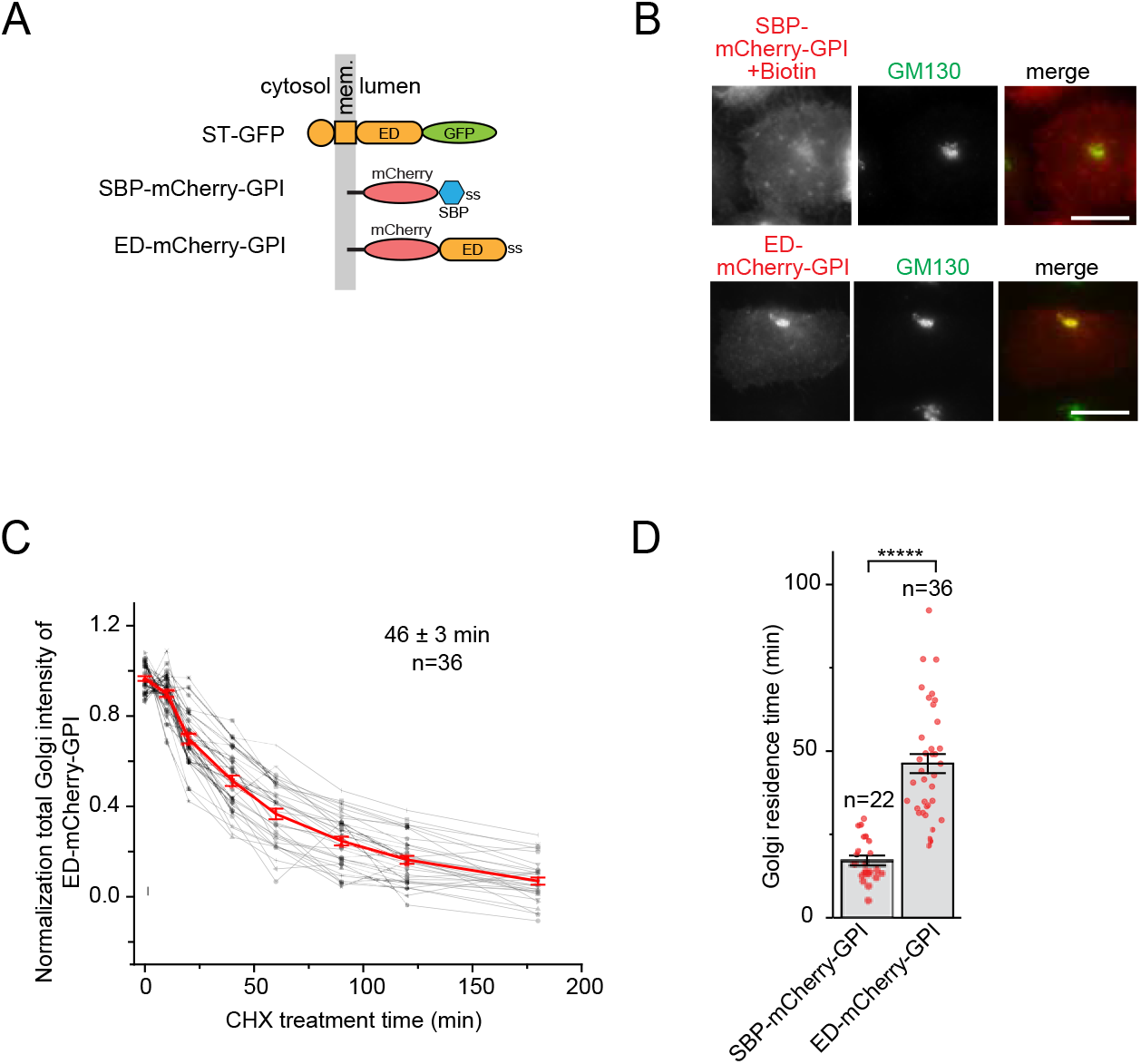
The ED of ST is sufficient for the Golgi retention. (A) The schematic diagram illustrating the domain organization of GPI-anchored chimeras. Mem., membrane; ss, signal sequence. (B) The GPI-anchored ED of ST localizes to the Golgi. HeLa cells transiently expressing indicated chimeras were immuno-stained for endogenous GM130. Cells expressing SBP-mCherry-GPI were incubated with 50 μM biotin for 20 h prior to immuno-staining. Scale bar, 20 μm. (C) The Golgi intensity of ED-mCherry-GPI vs time plot. HeLa cells transiently expressing ED-mCherry-GPI and GalT-mCherry were live imaged in the presence of CHX. (D) The Golgi residence times of ED-mCherry-GPI, which is from (C), and SBP-mCherry-GPI (control)(Sun et al., 2020). Error bar, mean ± standard error; *P* values are from *t* test (unpaired and two-tailed); *****, *P* ≤ 0.000005; n, the number of quantified cells.

## DISCUSSION

It is known that a small amount of a glycosyltransferase can escape the Golgi to the PM (Berger, 2002; Chen et al., 2000; Teasdale et al., 1994; Wong et al., 1992). Here, we elucidated the post-Golgi trafficking of glycosyltransferases. A glycosyltransferase such as ST is stably retained at the Golgi, with a Golgi residence time of ~ 5 h. It can very slowly exit the Golgi and follow the constitutive secretory pathway to reach the PM. At the PM, it is endocytosed to the endolysosome. Our observation indicated that a glycosyltransferase is mainly degraded by the cell surface ED-shedding instead of the lysosomal degradation, resulting in the release of its catalytic domain to the extracellular medium. Importantly, we demonstrated that most glycosyltransferases that we screened (11 out of 15) are not retrieved to the Golgi from the PM and endolysosome, suggesting that retention but not retrieval should be the primary mechanism for their Golgi localization.

The Golgi residence time was used as a metric for the Golgi retention of a glycosyltransferase as it is independent of the post-Golgi fate of the glycosyltransferase and can be conveniently measured by live cell imaging. By constructing many chimeras or mutants, we systematically and quantitatively assessed roles of the NCT, TMD and ED in the Golgi retention of ST. Our data are in agreement with what is known in the field that all regions are necessary and sufficient for the efficient Golgi retention of a glycosyltransferase (Banfield, 2011). We also showed that the TMD plays a significant role in the Golgi retention (Munro, 1991; Nilsson et al., 1991; Tang et al., 1992; Teasdale et al., 1992; Wong et al., 1992). Increasing the TMD length of ST was observed to reduce its Golgi residence time, consistent with the TMD-based sorting model (Bretscher and Munro, 1993; Munro, 1995a; Munro, 1995b).

Our systematic and quantitative study also yielded novel discoveries. We found that each of the three regions is sufficient to provide a Golgi retention. The retention effect of the three regions is additive and a synergistic effect between the NCT and TMD was revealed by our quantitative studies. Our Golgi residence time data indicate that the NCT length negatively affects the Golgi retention. Like a glycan’s role in the Golgi export (Sun et al., 2020), we think that the effect is likely due to physico-chemical properties of the NCT and their environment. Glycosyltransferases reside within the interior of the Golgi stack (Tie et al., 2018) and their NCTs are likely exposed to a tight inter-cisternal space (Engel et al., 2015). It is possible that large NCT is less compatible with the crowded molecular environment due to the steric hindrance (Stachowiak et al., 2013). On the other hand, the constitutive exocytic transport carrier generated at the cisternal rim might prefer transmembrane cargos with large NCTs, therefore carrying them out of the Golgi. At last, we showed that the GPI-anchored ED of ST strongly localizes to the Golgi. Different from the NCT and TMD, the Golgi retention of the ED is probably attributed to its interaction with endogenous glycosyltransferases within the cisternal interior (Kellokumpu et al., 2016).

Based on our data, we propose that long NCT and TMD can function as Golgi export signals for type II transmembrane cargos. Together with glycans (Sun et al., 2020), we have identified three types of Golgi export signals. A caveat of our and previous similar studies is the usage of overexpressed glycosyltransferases. Necessary kin-recognition partners are not present in stoichiometry in overexpressing cells. Hence, caution must be exercised in interpreting these findings, as it is unknown how faithful an overexpressed glycosyltransferase reflects the endogenous one in terms of membrane trafficking. The concern might be addressed in the future by testing genome-edited glycosyltransferases.

## MATERIALS AND METHODS

### DNA plasmids

See Table S1. The following DNA plasmids were previously described. GalT-mCherry (Lu et al., 2009), ST-GFP (Sun et al., 2020), B4GALT3-Myc (Tie et al., 2018), GL2 shRNA in pLKO.1 (Shi et al., 2018), pDMyc-Neo-N1 (Tie et al., 2018), Man1B1-Myc (Tie et al., 2018), SBP-mCherry-GPI (Boncompain et al., 2012), and TNFα-SBP-GFP (Boncompain et al., 2012). Human Genome Organization Gene Nomenclature Committee (HGNC) and Mouse Genome Informatics (MGI) official names of glycosyltransferases are used in this study except ST (ST6GAL1) and GalT (B4GALT1).

### Antibodies and small molecules

Mouse anti-EEA1 monoclonal antibody (mAb) (#610456; 1:500 for immunofluorescence or IF) and mouse anti-GM130 mAb (#610823; 1:500 for IF) were from BD Biosciences. Mouse anti-Lamp1 mAb (H4A3; 1:500 for IF) was from Developmental Studies Hybridoma Bank. Mouse anti-Myc mAb (#sc-40; 1:200 for IF), mouse anti-GAPDH mAb (#sc-47724; 1:1000 for Western blot) and mouse anti-GFP mAb (#sc-25778; 1:1000 for Western blot) were from Santa Cruz. Rabbit anti-GFP (Mahajan et al., 2019) and anti-mCherry (Sun et al., 2020) polyclonal sera were previously described. Horseradish peroxidase - conjugated goat secondary antibodies were from Bio-Rad. Alexa Fluor-conjugated goat antibodies against mouse or rabbit IgG (1:500 for IF) were from Thermo Fisher Scientific. CHX (working concentration: 10 μg/ml) and chloroquine (working concentration: 50 μM) were from Sigma Aldrich. Bafilomycin A1 (working concentration: 100 nM) was from Chemscene.

### Cell culture and transfection

HeLa and 293FT (Thermo Fisher Scientific) cells were cultured in Dulbecco’s Modified Eagle’s Medium supplemented with 10 % fetal bovine serum as previously described (Sun et al., 2020). HeLa cells were cultured on Φ 12 mm glass coverslips for fixed-cell imaging and Φ 35 mm glass bottom Petri dishes for live cell imaging as previously described (Sun et al., 2020).

### Lentivirus-transduced knockdown by shRNA

shRNA targeting GL2 (control), *VPS35, GOLPH3* or *COG4* cloned in pLKO.1 vector was transiently transfected to 293FT cells together with pLP1, pLP2, pLP/VSVG DNA plasmids (Thermo Fisher Scientific). Cell culture medium containing lentivirus particles was collected after 48 and 72 h. Virus enriched media were pooled, filtered through 0.45 μm filter (Sartorius) and used to infect HeLa cells in the presence of 8 μg/ml hexadimethrine bromide (Sigma-Aldrich, #H9268). After 24 h, the infection was repeated one more time. 24 h later, cells were transiently co-transfected with ST-GFP and GalT-mCherry. Live cell imaging was conducted in the presence of CHX to acquire the Golgi residence time. The knockdown efficiency was quantified by quantitative reverse transcription PCR (RT-qPCR). The total RNA was extracted using TRIzol™ reagent (Thermo Fisher Scientific). The subsequent reverse transcription was conducted using nanoScript 2 Reverse Transcription kit and random nanomer primers (Primerdesign). The real time PCR was performed on Bio-Rad CFX96 Touch using SYBR-green-based PrecisionFAST with LOW ROX qPCR kit (Primerdesign). The reading was normalized by that of β-tubulin transcript (*TUBB*). Below are primer pairs used: *TUBB* (5’-TTG GCC AGA TCT TTA GAC CAG ACA AC-3’ and 5’-CCG TAC CAC ATC CAG GAC AGA ATC-3’); *VPS35* (5’-AGT CGC CAT GCC TAC AAC ACA G-3’ and 5’-AGC TTG TTT TTG TCC AGG CAT C-3’); *GOLPH3* (5’-AGG ACC GCG AGG GTT ACA CAT C-3’ and 5’-ACA TCC CCT GTT GGA GCA TCT G-3’); *COG4* (5’-ACC GAA TGG GTC CTA ATC TGC AG-3’ and 5’-TGA TAG AGG CGG TTC TTG GCC AG-3’).

### Immunoprecipitation of the ED from the cell culture medium

HeLa cells were transiently transfected with ST-GFP. Next day, cells were incubated with 5 μg/ml VHH-mCherry for 1 h on ice to label cell surface ST-GFP. After extensive washing on ice, fresh culture medium was applied. Cells were subsequently warmed up to 37 °C for 0, 0.5, 1 and 2 h. The medium was collected and supplied with 50 mM HEPES and 2 mM DTT and cells were lysed in a buffer that has 50 mM HEPES, 100 mM NaCl, 1% Triton X-100 and 2 mM DTT. The cell culture medium was subjected to immunoprecipitation as described below. It was first incubated with 1 μl rabbit anti-mCherry polyclonal serum overnight at 0 °C. The system was then incubated with 20 μl protein A/G beads at 0 °C for 2 h. The Protein A/G beads together with captured antibody complex were pelleted and washed extensively. The bound proteins were eluted by boiling in 2 × SDS-sample buffer and subjected to Western blot analysis.

### Western blot

Proteins separated by the SDS-PAGE were transferred to polyvinyl difluoride membrane (Bio-Rad). The membrane was subsequently incubated with primary antibody and washed. After incubation with horse radish peroxidase-conjugated secondary antibody and extensive washing, the signal on the membrane was detected by chemiluminescence using LAS-4000 (GE Healthcare Life Sciences).

### VHH-mCherry purification and internalization assay

They were performed as previously described (Sun et al., 2020). Two types of internalization assays were performed: continuous and pulse-chase. Briefly, in the continuous internalization assay, 6×His-tagged VHH-mCherry purified from BL12DE3 *E Coli* cells or anti-Myc antibody was continuously incubated with live HeLa cells transiently expressing GFP or Myc-tagged reporter for indicated length of time at 37 °C. In the pulse-chase internalization assay, live HeLa cells transiently expressing GFP-tagged reporter were surface-labeled by VHH-mCherry on ice for 1 h. After washing, cells were warmed up to 37 °C for indicated length of time. After the internalization assay, cells were fixed and immuno-stained for indicated proteins.

### IF, microscopy, Golgi residence time and Golgi-to-cell intensity ratio

They were performed as previously described (Sun et al., 2020). Briefly, images were acquired under Olympus IX83 inverted microscope equipped with 63×/NA1.40 and 40×/NA1.20 objectives, 37 °C environment chamber, motorized filter cubes, focus drift correction device (ZDC2; Olympus), scientific complementary metal oxide semiconductor (Neo; Andor) and 200W metal-halide excitation light source (Lumen Pro 200; Prior Scientific). To measure the Golgi residence time, HeLa cells transiently coexpressing GFP-tagged reporter and GalT-mCherry were live-imaged under the treatment of CHX until the Golgi fluorescence almost disappeared. The resulting 2D-time lapse was segmented according to GalT-mCherry in ImageJ (https://imagej.nih.gov/ij/). The GFP fluorescence intensity within the Golgi was quantified and fitted to the first order exponential decay function *y*=*y_0_* + *A_1_* exp(-(*x-x_0_*)/*t_1_*) in OriginPro8.5 (OriginLab). The Golgi residence time was calculated as 0.693\**tl*. Time lapses with adjusted *R^2^* ≥ 0.80 and the length of acquisition ≥ 1.33\**t_1/2_* were considered.

## Data availability

All data are included in the manuscript and the Supporting Information.

## Author contributions

L.L. conceived and supervised the study. L.L. and X.S. designed the experiments. X.S. and B.C performed experiments. X.S., B.C., L.L. and Z.S. analyzed the data. L.L. and X.S. wrote the manuscript.

## Funding and additional information

This work was supported by the following grants to L.L.: MOE AcRF Tier1 RG35/17, Tier2 MOE2015-T2-2-073 and Tier2 MOE2018-T2-2-026.

## Abbreviations

The abbreviations used are:

AA: amino acid
CHX: cycloheximide
COG: conserved oligomeric Golgi
ER: endoplasmic reticulum
GalT: B4GALT1
GPI: glycosylphosphatidylinositol
IF: immunofluorescence
IPTG: isopropyl β-D-thiogalactopyranoside
mAb: monoclonal antibody
NCT: N-terminal cytosolic tail
PM: plasma membrane
ROI: region of interest
ST: ST6GAL1
TfR: transferrin receptor
TGN: *trans*-Golgi network
TMD: transmembrane domain
TNFα: tumor necrosis factor α
VHH: variable heavy-chain domain of heavy-chain-only antibody
VSVG: vesicular stomatitis virus glycoprotein G.

## Acknowledgements

We would like to thank the following researchers for sharing their DNA plasmids with us: R. Irvine, T. Kirchhausen, F. Perez, D. Root, E. Snapp and D. Stephens.

## SUPPLEMENTARY FIGURE LEGENDS

**Supplementary Figure 1.**
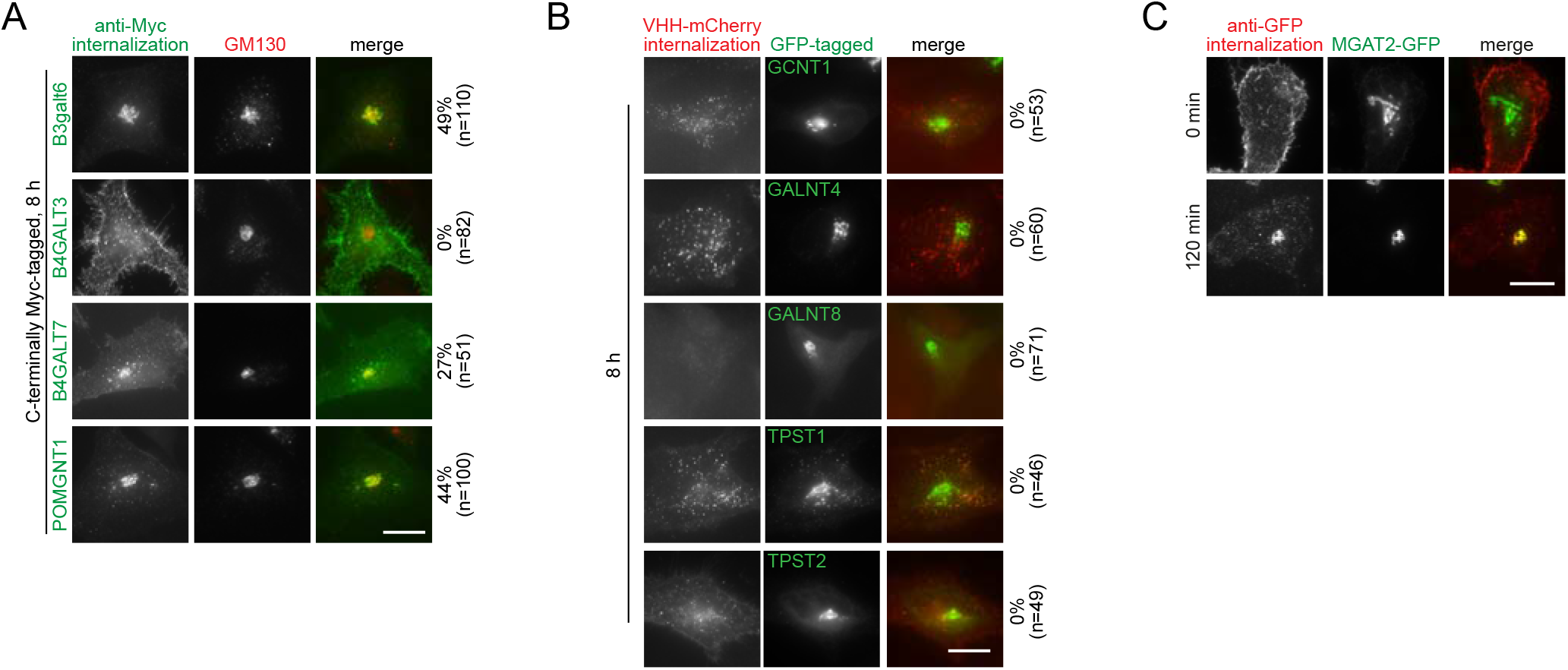
The endocytic trafficking of cell surface Golgi glycosyltransferases. (A) Cell surface B3galt6, B4GALT7 and POMGNT1, but not B4GALT3, can be retrieved to the Golgi, although to a very limited extent. HeLa cells transiently expressing indicated glycosyltransferases were continuously incubated with anti-Myc antibody for 8 h at 37 °C before immuno-staining for Myc-tag and endogenous GM130. (B) Cell surface GCNT1, GALNT4, GALNT8, TPST1 and TPST2 are not retrieved to the Golgi. HeLa cells transiently expressing indicated glycosyltransferases were continuously incubated with VHH-mCherry for 8 h at 37 °C before imaging. Note that the surface pool of GALNT8-GFP was undetectable by our assay. (C) The retrieval of cell surface MGAT2-GFP to the Golgi as monitored by anti-GFP serum. HeLa cells transiently expressing MGAT2-GFP were surface-labeled with rabbit polyclonal anti-GFP serum on ice. After washing, cells were warmed up to 37 °C for indicated time and immuno-stained for rabbit IgG. Scale bar, 20 μm. The number on the right of each panel indicates the percentage of cells displaying the colocalization of anti-Myc antibody (A), VHH-mCherry (B) or anti-GFP antibody (C) with the Golgi; n, the number of cells counted.

**Supplementary Figure 2.**
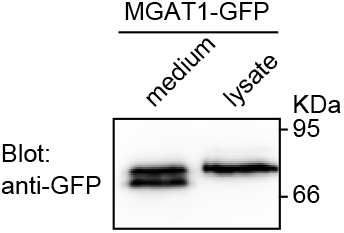
The ED of MGAT1-GFP can be detected in the culture medium. The experiment was conducted as described in Figure 2A. Molecular weights (in KDa) are labeled at right.

**Supplementary Figure 3.**
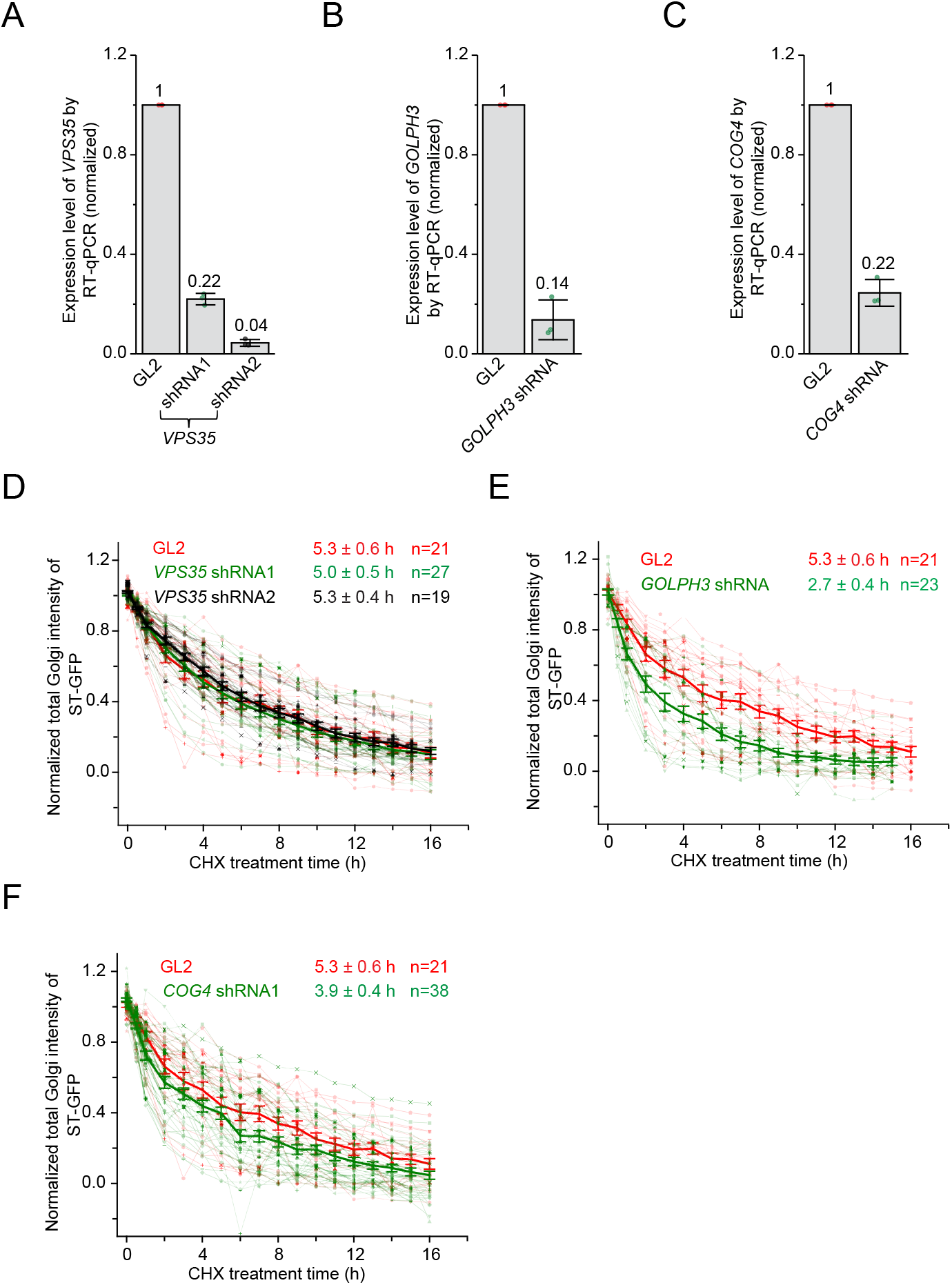
The knockdown of *VPS35, GOLPH3* and *COG4* and measuring the Golgi residence times of ST-GFP in the resulting knockdown cells. This figure corresponds to Figure 3, B-D. (A-C) Expression levels of *VPS35, GOLPH3* and *COG4* were knocked down by corresponding shRNAs. HeLa cells were subjected to lentivirus-mediated transduction of indicated shRNAs and mRNA levels of target genes were quantified by RT-qPCR after two days. The expression data were normalized by that of GL2 control shRNA. Error bar, mean ± standard deviation; three independent experiments were conducted. (D-F) Acquiring Golgi residences times of ST-GFP in knockdown cells. HeLa cells were subjected to lentivirus-mediated transduction of indicated shRNAs. After two days, cells were transiently transfected to express ST-GFP and GalT-mCherry. In the next day, live cell imaging was performed under the treatment of CHX. The total fluorescence intensity within the Golgi (marked by GalT-mCherry) was quantified at each time point, normalized by that of 0 h and subsequently plotted against the time. Each intensity series was fitted to the first order exponential decay function to calculate the Golgi residence time (indicated). Individual and mean time series are represented by faded and solid color lines, respectively. Error bar, mean ± standard error; n, the number of quantified cells.

**Supplementary Figure 4.**
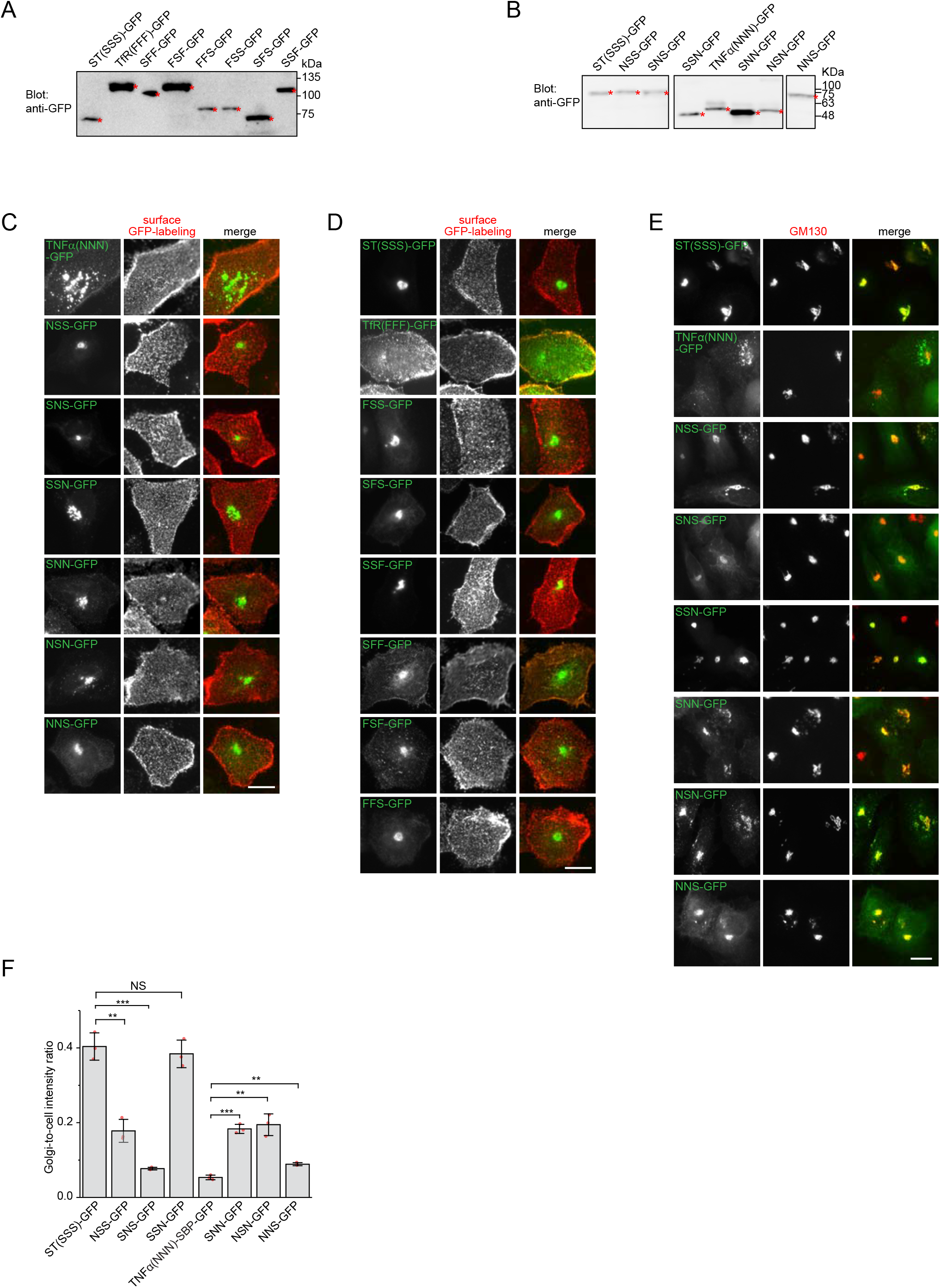
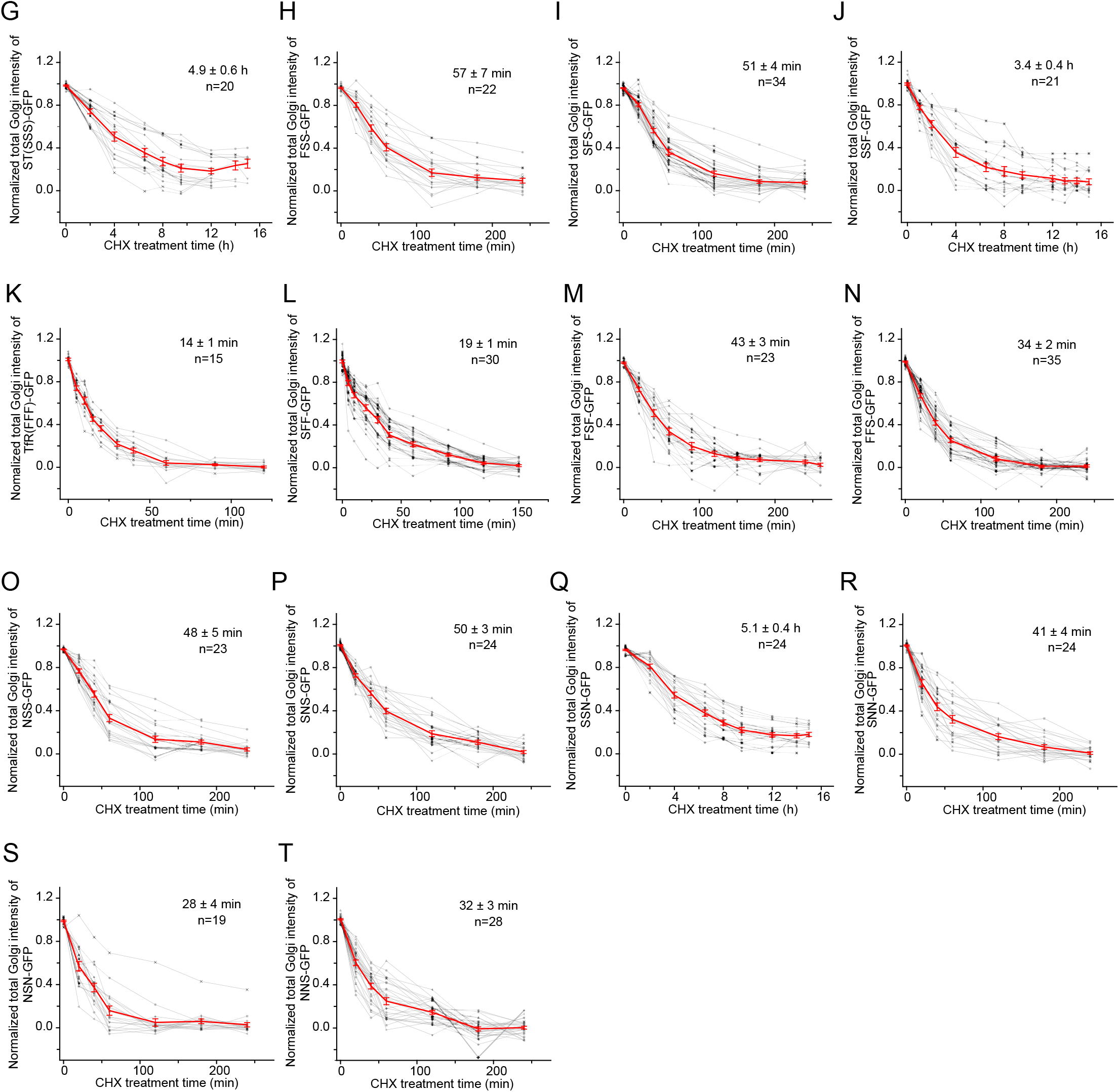
Western blots, subcellular localization and the Golgi intensity vs time plots of ST swapping chimeras. This figure corresponds to Figure 4. (A,B) ST swapping chimeras shown in Figure 4, A and B, express proteins of expected sizes in Western blot. Lysates of HeLa cells transiently expressing GFP-tagged chimeras were subjected to Western blot analysis using anti-GFP antibody. Swapping chimeras between ST and TfR are in (A) while those between ST and TNFα are in (B). * indicates correct bands. Molecular weights (in KDa) are labeled at right. (C,D) Positive surface staining of ST swapping chimeras shown in Figure 4, A and B, demonstrates that their C-termini are exposed to the extracellular space, as expected for type II transmembrane proteins. HeLa cells transiently expressing indicated C-terminally GFP-tagged chimeras were surface-labeled with rabbit polyclonal anti-GFP serum. Cells were subsequently immunostained for rabbit IgG and imaged. (E) The subcellular localization of swapping chimeras between ST and TNFα. HeLa cells transiently expressing swapping chimeras were immuno-stained for endogenous GM130. Scale bar, 20 μm. (F) The Golgi-to-cell intensity ratios of swapping chimeras between ST and TNFα. Quantification was performed on cells described in (E). Error bar, mean ± standard deviation; n= 3 independent experiments were conducted with ≥ 24 cells quantified in each experiment. *P* values are from *t* test (unpaired and two-tailed); NS, not significant; **, *P* ≤ 0.005; ***, *P* ≤ 0.0005. (G-T) The Golgi intensity vs time plots of ST chimeras under CHX treatment. The experiment corresponds to Figure, 4 E and F. The total fluorescence intensity within the Golgi (marked by GalT-mCherry) was quantified at each time point, normalized and subsequently plotted against the time. Each intensity series was fitted to the first order exponential decay function to calculate the Golgi residence time (indicated). Individual and mean time series are represented by grey and red color lines, respectively. Error bar, mean ± standard deviation (F) or mean ± standard error (G-T); n, the number of quantified cells.

**Supplementary Figure 5.**
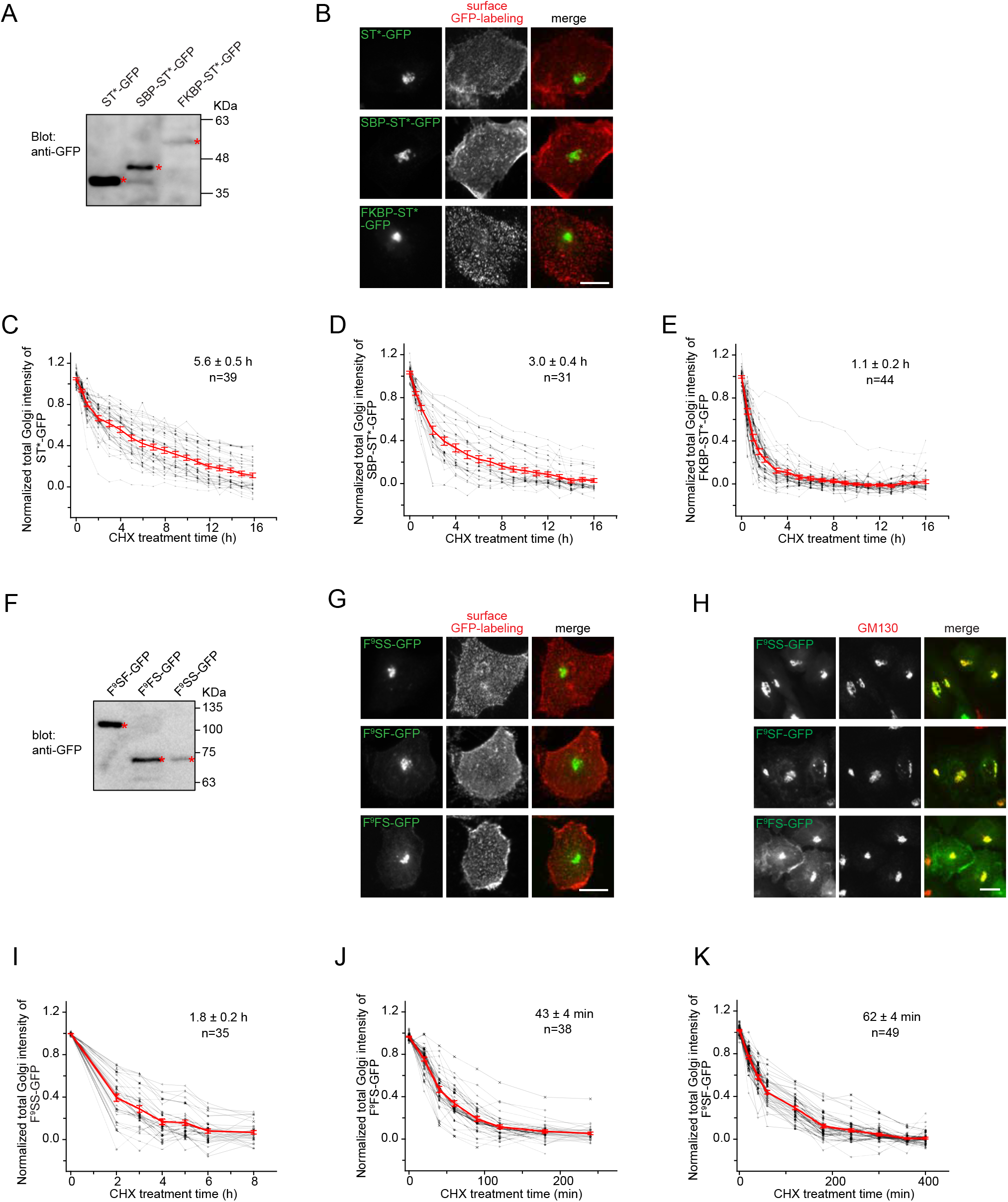
Western blots, subcellular location and the Golgi intensity vs time plots of chimeras with different NCT lengths. This figure corresponds to Figure 5. (A) ST* chimeras shown in Figure 5A express proteins of expected sizes in Western blot. Lysates of HeLa cells transiently expressing GFP-tagged ST* chimeras were subjected to Western blot analysis using anti-GFP antibody. * indicates correct bands. (B) Positive surface staining of ST* chimeras shown in Figure 5A demonstrates that their C-termini are exposed to the extracellular space, as expected for type II transmembrane proteins. The experiment was conducted as described in Figure S4, C and D. (C-E) The Golgi intensity vs time plots of ST* chimeras shown in Figure 5A. The experiment corresponds to Figure 5B. The data are processed as described in Figure S4, G-T. (F,G) C-terminally-GFP tagged F^9^SF, F^9^FS and F^9^SS express proteins of expected sizes in Western blot and were positively surface stained by anti-GFP antiserum. The experiments were similar to (A,B). (H) The subcellular localization of C-terminally-GFP tagged F^9^SF, F^9^FS and F^9^SS. HeLa cells transiently expressing indicated chimera were immuno-stained for endogenous GM130. (I-K) Golgi intensity vs time plots of C-terminally-GFP tagged F^9^SF, F^9^FS and F^9^SS. The experiment corresponds to Figure 5, G-I and the data were processed as described in Figure S4, G-T. In (A,F), molecular weights (in KDa) are labeled at right. Scale bar, 20 μm.

**Supplementary Figure 6.**
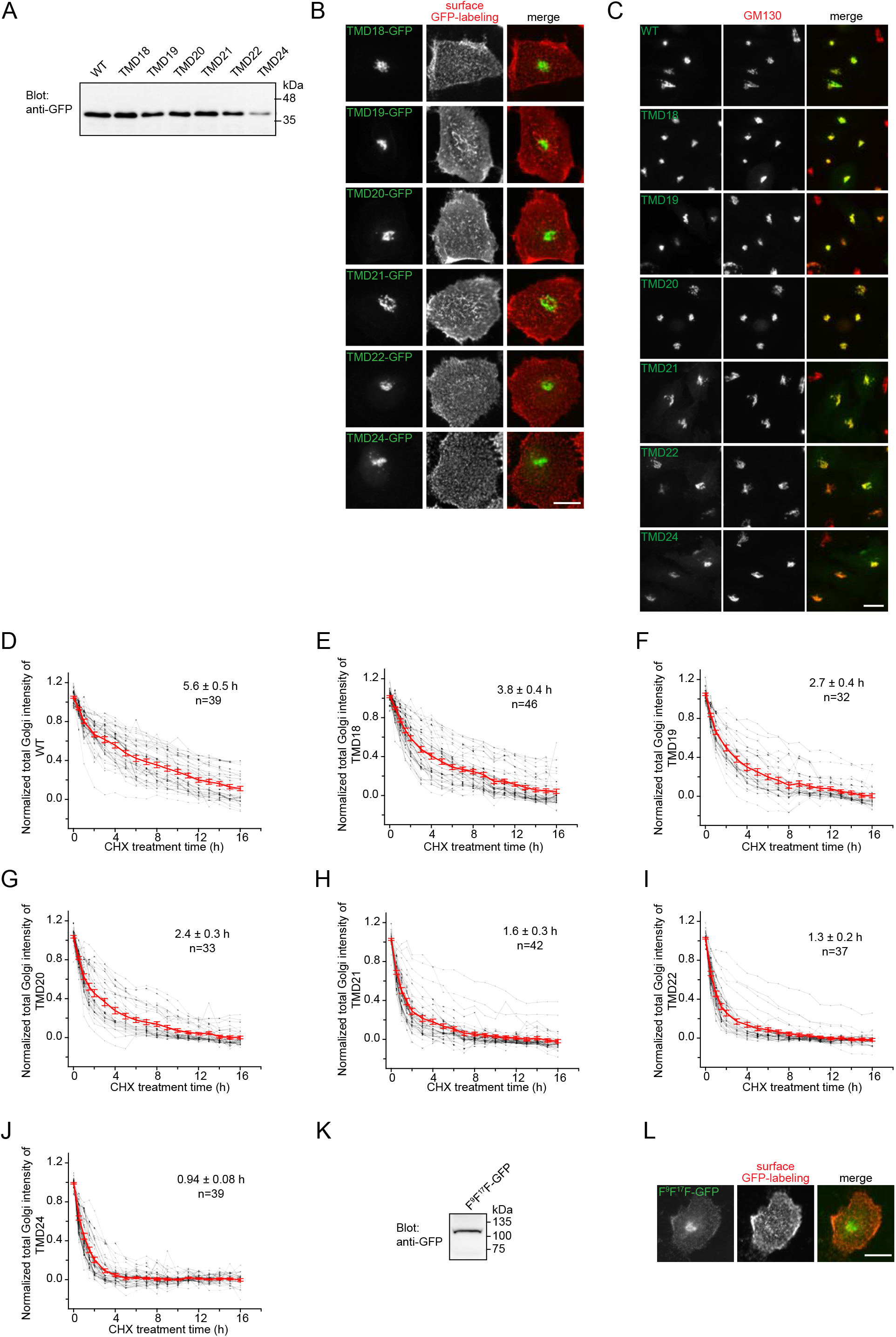
Western blots, surface staining, subcellular localization and the Golgi intensity vs time plots of ST* mutants with increasing TMD lengths. (A) TMD mutants of ST* shown in Figure 6A express proteins of expected sizes. Lysates of HeLa cells transiently expressing GFP-tagged TMD mutant of ST* were subjected to Western blot analysis using anti-GFP antibody. (B) Positive surface staining of ST* TMD mutants shown in Figure 6A demonstrates that their C-termini are exposed to the extracellular space, as expected for type II transmembrane proteins. The experiment was conducted as described in Figure S4, C and D. (C) ST* TMD mutants shown in Figure 6A localize to the Golgi. HeLa cells transiently expressing indicated ST* TMD mutant were immuno-stained for endogenous GM130. (D-J) The Golgi intensity vs time plots of ST* TMD mutants. The experiment was conducted and resulting data were processed as described in Figure S4, G-T. (K,L) F^9^F^17^F-GFP expresses a protein of expected size and it displays positive surface staining, as expected for a type II transmembrane protein. Experiments were conducted similar to those in (A,B). In (A,K), molecular weights (in KDa) are labeled at right. In (B, C, L), scale bar, 20 μm.

**Supplementary Figure 7.**
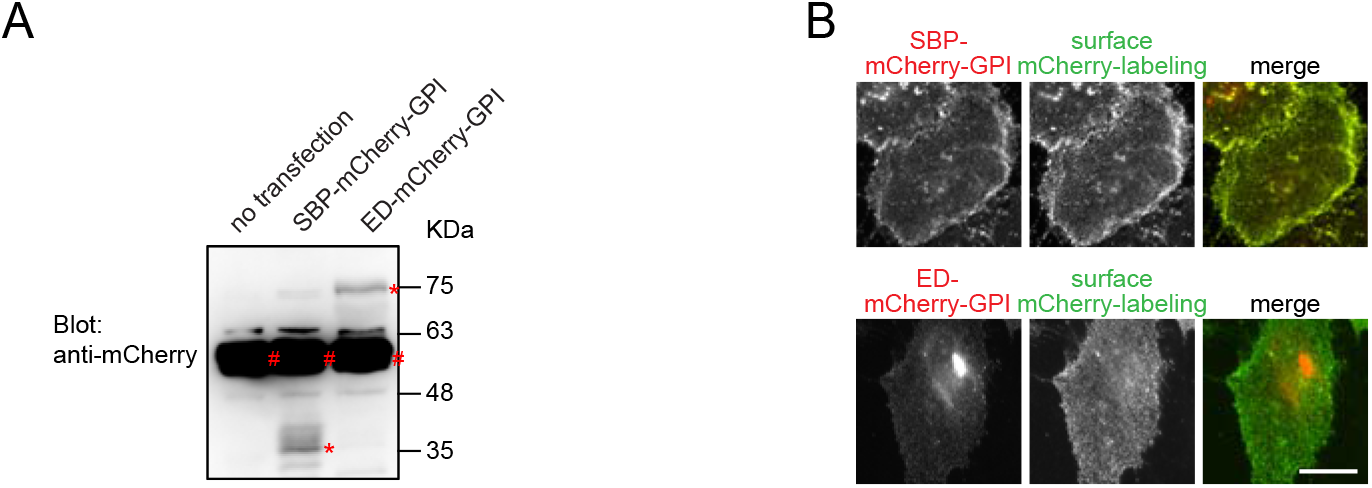
Western blot and cell surface staining of GPI-anchored chimeras. (A) ED-mCherry-GPI and SBP-mCherry-GPI constructs express proteins of expected sizes. Lysates of HeLa cells transiently expressing indicated proteins were subjected to Western blot analysis using anti-mCherry antibody. *, specific band; #, non-specific band. Molecular weights (in KDa) are labeled at right. (B) Positive surface staining of ED-mCherry-GPI and SBP-mCherry-GPI demonstrates that their mCherry groups are exposed to the extracellular space, as expected for GPI-anchored proteins. The experiment was conducted as described in Figure S4, C and D. Cells expressing SBP-mCherry-GPI were incubated with 50 μM biotin for 20 h prior to surface staining. Scale bar, 20 μm.

**Table S1.**
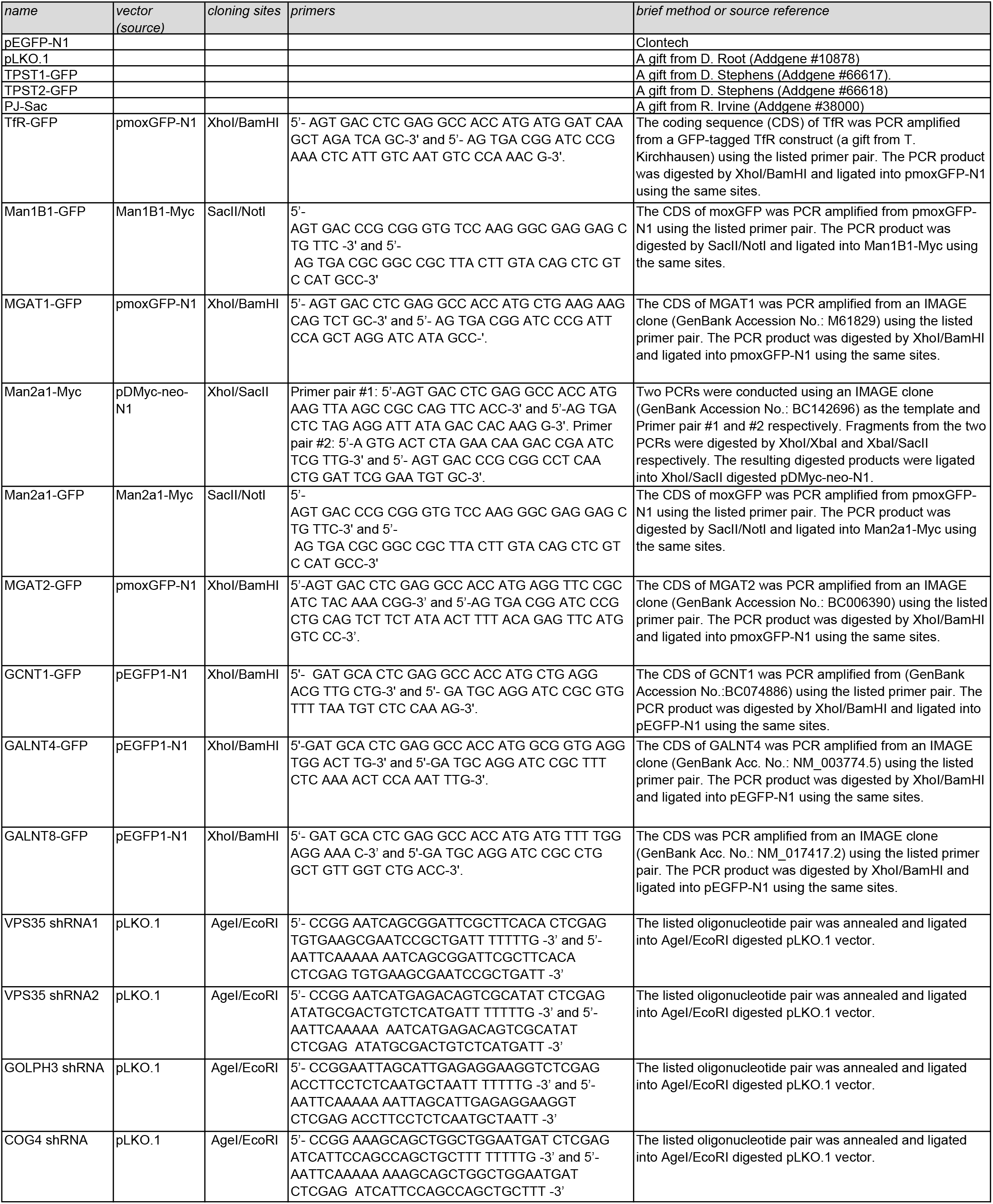
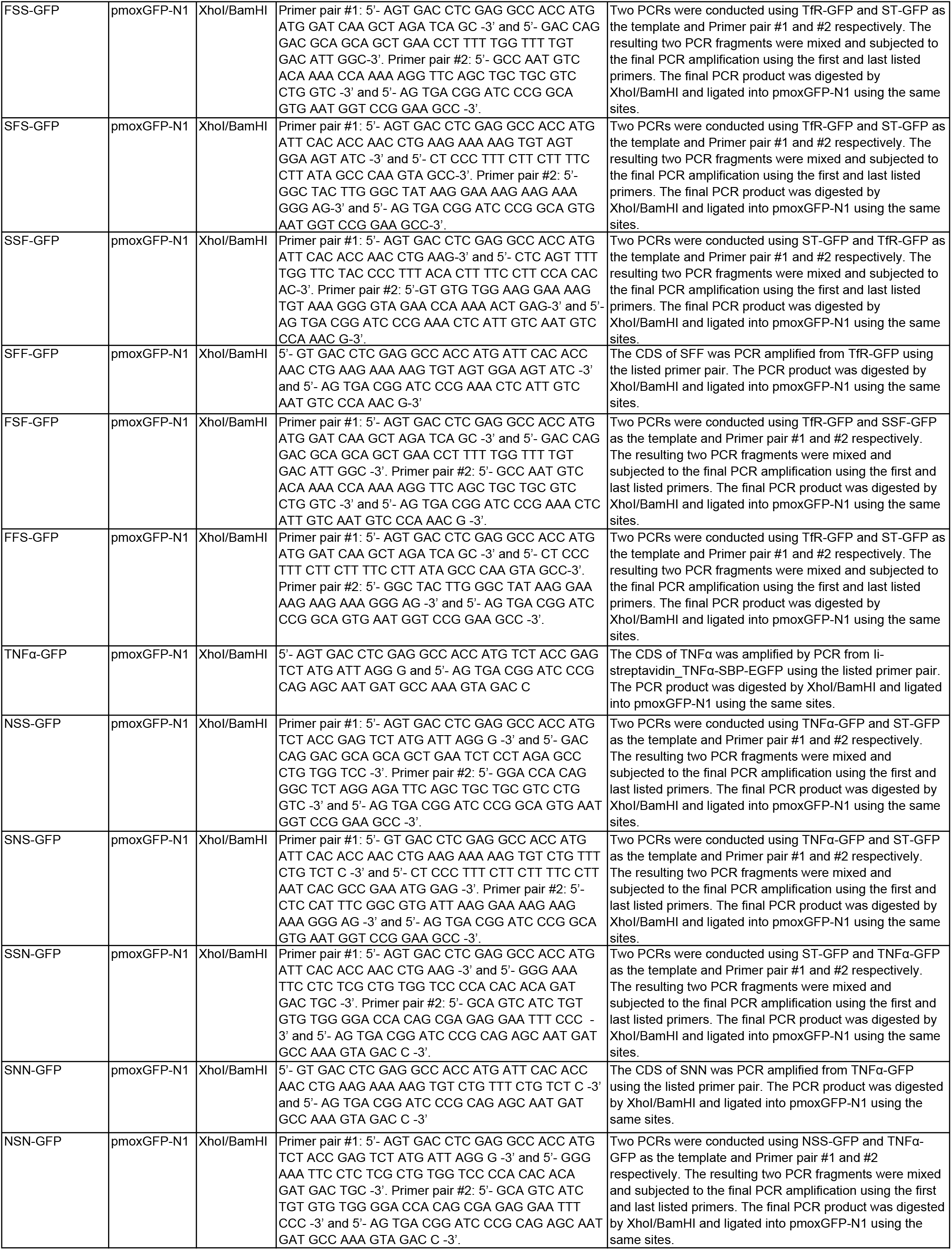
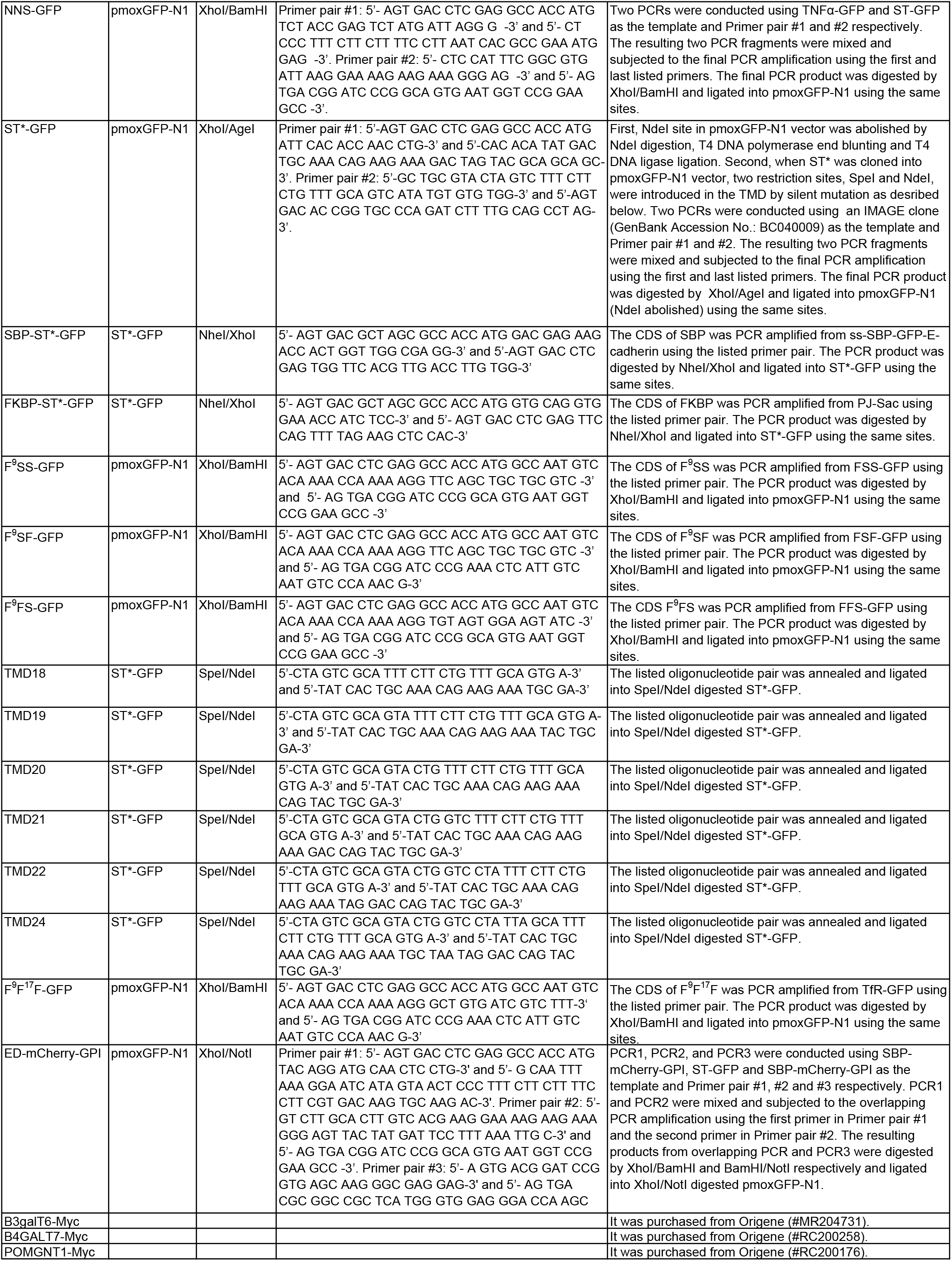
List of DNA plasmids.

